# The convergent evolution of hummingbird pollination results in repeated floral scent loss through gene downregulation

**DOI:** 10.1101/2024.05.08.592797

**Authors:** Kathy Darragh, Kathleen M. Kay, Santiago R. Ramírez

## Abstract

The repeated evolution of the same trait in different lineages provides powerful natural experiments to study the phenotypic and genotypic predictability of how traits are gained and lost. A fascinating example of this is the repeated evolution of hummingbird pollination in plant lineages in the Americas, a widespread and often unidirectional phenomenon. The spiral gingers in the genus *Costus* are ancestrally bee-pollinated, and hummingbird pollination has evolved multiple times independently in the tropical Americas. These pollinator transitions are accompanied by predictable morphological and color changes, but the changes in floral scent have not been described. In this study, we describe the floral scent composition of 30 species of *Costus* sampled across the phylogeny to understand how floral scent has evolved across the genus with respect to pollinator transitions. We then combine transcriptomics and genomics to identify genetic expression differences and gene family evolution associated with pollinator transitions. We show that hummingbird-pollinated species have mostly lost their floral scent, whereas bee-pollinated species exhibit either floral scent maintenance or in some cases, gains of more diverse scent profiles. We find the floral scent loss appears to be due to gene downregulation rather than pseudogenization. The remarkable consistency of scent loss in hummingbird-pollinated species highlights the shared strong selection pressures experienced by these lineages. Even species with more recent transitions from bee to hummingbird pollination exhibit scent loss, highlighting the rapid breakdown of scent production following pollinator transitions. This research highlights the capacity for rapid changes when selection pressures are strong through downregulation of floral scent genes.

**Resumen:** La evolución independiente de un rasgo en diferentes linajes ofrece la oportunidad de estudiar la predictibilidad de fenotipos y sus bases genéticas. Un ejemplo fascinante es la evolución repetida de polinización por colibríes en varios linajes de plantas en las Américas, un fenómeno generalizado y típicamente unidireccional. Las especies de caña agria del género *Costus* son polinizadas ancestralmente por abejas y la polinización por colibríes ha evolucionado varias veces de forma independiente en América tropical. Cada transición de polinizador ha sido acompañada de varios cambios morfológicos y colores predecibles, pero los cambios en el aroma floral no han sido descritos. En este estudio describimos la composición del aroma floral de 30 especies de *Costus* para comprender cómo el aroma floral ha evolucionado con respecto a las transiciones de polinizadores. Combinamos transcriptómica y genómica en un clado de *Costus* para identificar diferencias en la expresión genética y la evolución de la familia de genes asociadas con las transiciones de polinizadores. Nuestros resultados muestran que las especies polinizadas por colibríes han perdido en su mayoría su aroma floral, mientras que las especies polinizadas por abejas ya sea mantienen el aroma floral y, en algunos casos, muestran ganancias de perfiles de aroma más diversos. La consistencia en la pérdida de olor en especies polinizadas por colibríes resalta las fuertes presiones de selección natural que experimentan estos linajes. Incluso las especies con transiciones más recientes de polinización de abejas a colibríes exhiben pérdida de olor, lo que destaca la rápida interrupción de la producción de olores después de las transiciones de los polinizadores. También encontramos que la pérdida de aroma floral parece estar controlada por regulación génica negativa y no por la pérdida de genes de producción de aromas (*e.g.* pseudogenización). Esta investigación destaca la rapidez de cambios a través de la regulación negativa de genes que controlan aromas florales cuando las presiones de selección son fuertes.

**Significance Statement:** The independent evolution of the same trait in different lineages provides a natural experiment to investigate the predictability of evolution. A fascinating example is the evolution of hummingbird pollination, which has occurred many times across flowering plants, and often involves convergence in multiple traits. In this study, we combine comparative chemical ecology with genomics and transcriptomics to study floral scent evolution in the genus *Costus*. We show that species exhibit loss of floral scent following transitions to hummingbird pollination suggesting a shared strong selection pressure favoring scent loss. We demonstrate the floral scent loss appears to be due to gene downregulation. This research highlights the capacity for rapid changes when selection pressures are strong through downregulation of scent genes.

## Introduction

The repeated evolution of the same traits in different lineages is thought to reflect shared selection pressures. These cases of repeated evolution apply not only to the gain of traits, such as the evolution of antifreeze proteins in fish (Chen et al. 1997) or carnivory in plants (Albert et al. 1992), but also the loss of traits, as seen in the eye regression of cavefish (Sifuentes-Romero et al. 2023), or reduction of attractive traits such as flower size or floral scent in self-fertilizing plants (Tsuchimatsu and Fujii 2022). Repeated evolution provides us with systems where we can study the process of adaptation and ask questions about the predictability of evolution, and the extent of constraint, or flexibility, of the underlying molecular mechanisms (Orr 2005; Losos 2011; Sackton and Clark 2019).

One such example of repeated evolution is pollinator transitions in flowering plants. For example, the evolution of hummingbird pollination has occurred many times across flowering plants, and often involves convergence in a suite of traits. Flowers pollinated by hummingbirds are characterized by traits thought to attract and fit hummingbirds (such as a narrow shape and large amounts of nectar) and in addition, traits thought to deter bees (red coloration, lack of nectar guides, lack of scent) (Schemske and Bradshaw 1999; Castellanos et al. 2004; Bergamo et al. 2016). Interestingly, the evolution of hummingbird pollination has primarily been documented in one direction, from ancestral bee pollination to derived hummingbird pollination (Rosas-Guerrero et al. 2014; Abrahamczyk and Renner 2015; Kay and Grossenbacher 2022). However, there is clade-specific variation in this directionality, with the reasons for pollinator transitions remaining unclear (Tripp and Manos 2008; Rosas-Guerrero et al. 2014; Abrahamczyk and Renner 2015; Kay and Grossenbacher 2022; Barreto et al. 2024).

Hummingbird-pollinated flowers often lack floral scent and therefore provide an excellent system to study repeated trait loss (Knudsen et al. 2004; Wessinger 2024). Genomic studies have shown that, even for birds, hummingbirds have a low number of olfactory receptors (Driver and Balakrishnan 2021; Driver 2022), suggesting at least a reduced investment in this sensory modality. Behavioral experiments have also suggested that hummingbirds rely more on visual than olfactory cues when foraging (Goldsmith and Goldsmith 1982). In fact, floral scent analyses of hummingbird pollinated flowers have shown either complete lack of floral scent or low emission rates of floral scent compounds. (Knudsen et al. 2004; Byers et al. 2014; Amrad et al. 2016). The lack of floral scent in hummingbird-pollinated species suggested that there is a cost to producing scent for these species. This could be due to energetic costs of compound production, but this is predicted to be minimal (Raguso 2016; Pichersky and Raguso 2018). An alternative is that there is an ecological cost for hummingbird-pollinated species attracting bees, also called the bee avoidance hypothesis (Castellanos et al. 2004; Bergamo et al. 2016).

A fascinating group in which to study the evolution of hummingbird pollination and potential trait loss is the genus *Costus* (Costaceae). The genus contains over 80 species of perennial understory herbaceous monocots, the majority of which have diverged rapidly within the last 3 million years in Central and South America (Maas 1972; Maas 1977; Vargas et al. 2020). Since establishing in the American tropics, hummingbird pollination has evolved repeatedly and independently in *Costus* from an ancestral state of bee pollination (Vargas et al. 2020). Specifically, bee-pollinated species are pollinated primarily by orchid bees (Apidae: Euglossini), with females visiting flowers for nectar and pollen, and males visiting for nectar (Kay and Schemske 2003). Hummingbird-pollinated species are primarily pollinated by non-territorial hermit hummingbirds (Kay and Schemske 2003). While different species were initially classified as hummingbird or bee pollinated (Maas 1972; Maas 1977), many species do not fit the classic paradigm of long tubular red flowers associated with hummingbird pollination (Grant and Grant 1968). For example, many bee-pollinated flowers have some red coloration (Kay and Schemske 2003), and hummingbird-pollinated flowers are short (Yost and Kay 2009). Despite this variation, a single phenotypic optimum is shared across most of the independent transitions to hummingbird pollination in the clade (Kay and Grossenbacher 2022). Traits including small flower size, the absence of nectar guides, and brightly colored floral bracts are all predictive of hummingbird pollination (Kay and Grossenbacher 2022). Furthermore, *Costus* species show extensive sympatry between different pollination syndromes (Kay and Schemske 2003; Vargas et al. 2020; Kay and Grossenbacher 2022) and even the specialist herbivores are shared across *Costus* species (García-Robledo et al. 2013), allowing us to isolate the effects of pollination syndrome from geographic location or herbivory. The combination of recent genomic resources, a well-resolved phylogeny, documented pollinators, and the possibility of collecting samples both in the field and in greenhouses, makes *Costus* a powerful system to study the evolution of convergence on hummingbird pollination (Vargas et al. 2020; Kay and Grossenbacher 2022; Harenčár et al. 2023).

While the morphological and color changes involved in the evolution of hummingbird pollination have been well-studied in *Costus*, the expected scent changes have not been characterized. In this study, we describe floral scent in 30 species of *Costus* sampled across the phylogeny and including eight independent origins of hummingbird pollination. We test for differences in floral scent between bee-pollinated and hummingbird-pollinated species and investigate how floral scent has evolved across the genus. We then focus on one clade of *Costus* with variation in floral scent production and generate floral transcriptomic data. We use this data to identify gene expression differences associated with floral scent production to better understand the genetic changes underlying floral scent evolution in this group. Finally, we annotated a family of genes important for the production of terpenoid floral scent compounds (the terpene synthase genes (TPS)) in three previously available *Costus* genomes, and a high-quality genome assembly for *Costus allenii* generated in this study, encompassing two independent transitions to hummingbird pollination. High-quality annotations allow us to analyze the evidence of gene loss or pseudogenization following transition to hummingbird pollination in this gene family that plays an important role in floral scent evolution. Our study, therefore, combines multiple approaches to characterize repeated phenotypic evolution and the molecular mechanisms of these changes with shifts to hummingbird pollination in *Costus*.

## Results

### Bee-pollinated species produce more compounds and more diverse floral scents

We collected 100 samples from 30 *Costus* species including species previously described as both bee- and hummingbird-pollinated (Fig 1A, Table S1). Using multivariate techniques, we visualized the floral scent profiles to identify the broad patterns in our dataset. Before filtering our data, we found that samples from hummingbird-pollinated species did not differ from ambient control samples (Fig 1B). In contrast, samples from bee-pollinated species spread across a wider section of chemospace, with many samples visually separated from both ambient and hummingbird-pollinated samples (Fig. 1B). We found that both pollination group, and species (nested within pollination group) explained variation in floral scent (PERMANOVA, pollination group: F_2,167_=8.0, p<0.001; species: F_28,167_=2.5, p<0.001). Interestingly, bee-pollinated species were significantly different from both hummingbird-pollinated species and ambient control samples (Pairwise PERMANOVA, p=0.003, whereas hummingbird-pollinated species did not differ from ambient control samples (Pairwise PERMANOVA, p=0.07) (Figure S1).

**Figure 1.**
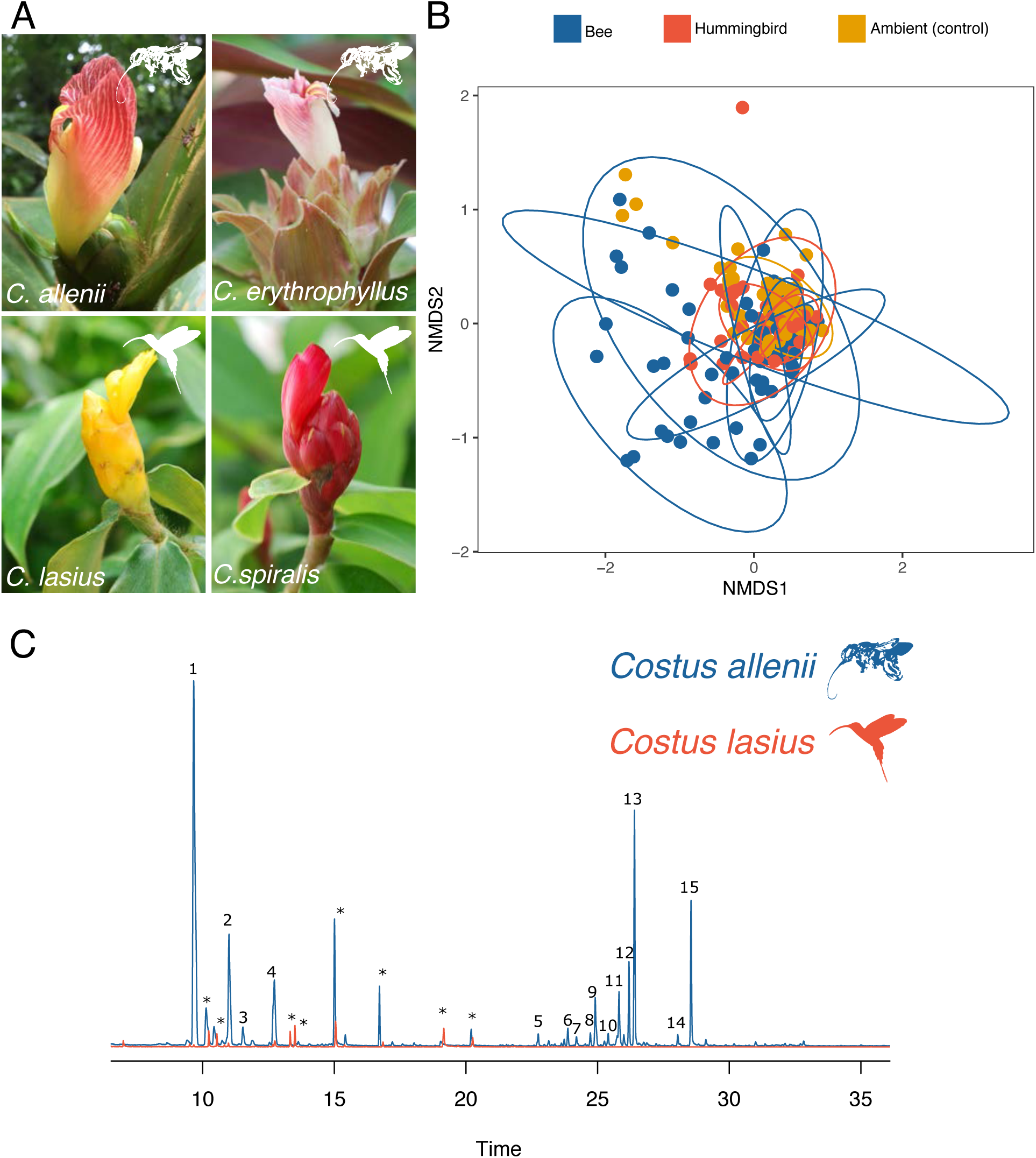
A) Representative images of two pairs of closely related bee- and hummingbird-pollinated species: *Costus allenii* and *C. lasius* and *C. erythrophyllus* and *C. spiralis*. B) NMDS (nonmetric multidimensional scaling) plot illustrating in the variation in the unfiltered dataset (Stress=0.12). Points are labelled as bee-pollinated, hummingbird-pollinated, or ambient samples. Ambient samples are control samples of the air taken in the same area as the focal plant individual. C) Total ion chromatogram of floral scent from *Costus allenii* and *C. lasius*. We chose the individuals with the second-highest amount of scent from each species to avoid plotting an outlier for either group. The *C. lasius* sample was shifted on the x-axis by 1.5 minutes to match the *C. allenii* sample due to changes in the GC/MS column over time. Abundance is scaled to the highest peak present in *C. allenii*. 1, alpha-pinene; 2, beta-pinene; 3, beta-myrcene; 4, cineole; 5, alpha-copaene; 6, (*E*)-beta-caryophyllene; 7, gamma-elemene; 8, alpha-humulene; 9, (*Z*)-beta-caryophyllene; 10, germacrene D; 11, alpha-elemene; 12, gamma-cadinene; 13, delta-cadinene; 14, germacrene-d-4-ol; 15, unidentified sesquiterpene (RI=1616). Contaminants are identified with an asterisk (*).

After filtering likely contaminants from the ambient air, we obtained a final dataset with 60 floral scent compounds (Table S2). The majority of these compounds were terpenoid (87%, 52/60), with a few aromatics and compounds of unknown class. The dataset included both monoterpenes and sesquiterpenes. No one compound class was restricted to a specific pollinator group. After removing the ambient compounds, we found that one bee-pollinated sample and eighteen hummingbird-pollinated samples had no detectable floral scent, confirming the similarity of the hummingbird-pollinated samples to the ambient air samples. The unscented hummingbird-pollinated samples were from six species. For two species, *C. montanus* and *C. stenophyllus*, all individuals produced no detectable floral scent (4/4 individuals). Furthermore, *most C. chartaceus* (4/5), *C. lasius* (4/7), and *C. wilsonii* (5/8), and a single *C. productus* individual (1/5) lacked floral scent.

We found that bee-pollinated flowers produced more floral scent compounds, averaging 9 (s.d. 10) compounds in comparison to only 2 (s.d. 4) for hummingbird-pollinated flowers (Figure 1C). Bee-pollinated flowers contained more compounds than hummingbird-pollinated flowers (Fig 2A). We confirmed that this was the case for both plants raised in a greenhouse and those sampled in the field (Fig. S2). Similar to the number of compounds, we used a measure of biochemical and structural diversity of the compounds present in a sample to calculate the overall chemical diversity. The Functional Hill Diversity index considers not only the total number of compounds but also their biosynthetic complexity. We found that the bee-pollinated floral scents had a higher chemical diversity (Fig 2B). Bee-pollinated flowers also produced a higher abundance of floral scent, as measured by total ion abundance in each sample, a proxy for compound amount (Fig. 2C).

**Figure 2.**
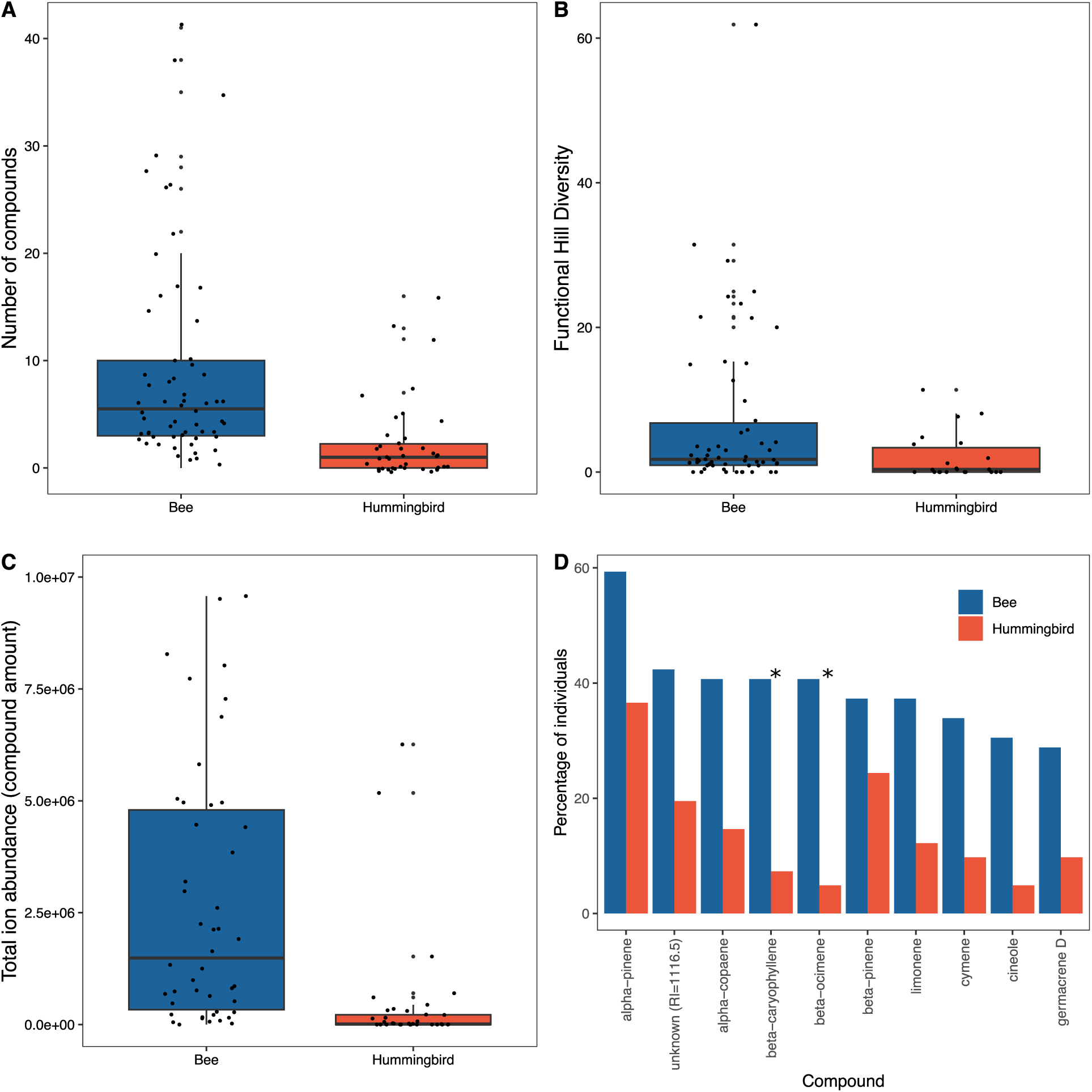
Differences in bee-pollinated and hummingbird-pollinated *Costus* species in terms of A) Number of scent compounds produced (ANOVA, pollination group: F_1,70_=60.4, p<0.0001, species: F_28,70_=9.4, p<0.0001), B) The diversity of scent compounds produced (ANOVA, pollination group: F_1,51_=5.3, p=0.03, species: F_27,51_=1.4, p=0.1), C) Total compound amount produced (measured by proxy of total ion abundance) (ANOVA, pollination group: F_1,70_=109, p=<0.0001, species: F_28,70_=2.8, p<0.001) and D) the percentage of individuals that produced the ten most common floral scent compounds in the dataset. Significant differences in frequency between bee- and hummingbird-pollinated flowers as determined by Chi-squared testing are highlighted with an asterisk. Fifteen individuals with the highest compound production were removed from C for visualization.

To see how individual compounds varied between bee- and hummingbird-pollinated species we calculated the percentage of individuals in each pollination group that produced the most common floral scent compounds in the dataset. We found that these compounds are present in a higher proportion of bee-pollinated flowers than hummingbird-pollinated flowers (Fig. 2D). In particular, (*E*)-beta-ocimene and beta-caryophyllene show the largest percentage differences between the two pollination groups (Fig. 2D) and are the only two compounds to exhibit a significant difference in frequency between pollinator groups (Table S2).

To facilitate visualization of our final filtered dataset using NMDS we removed the nineteen samples with no detectable floral scent. When this filtered dataset was plotted, we observed more overlap between the hummingbird- and bee-pollinated samples, but the two sets of samples significantly different (Figure S1).

### Phylogenetic analyses show that hummingbird-pollinated species have lost floral scent

To understand the evolution of floral scent in *Costus*, we carried out phylogenetic analyses using a previously published phylogeny (Vargas et al. 2020). In our dataset, we found a strong correlation between pollination group and the phylogenetic covariance matrix using two-block partial least squares analysis (rPLS = 0.56; P=0.01). This reduces the statistical power of testing group differences using phylogenetic ANOVA due to the high aggregation of pollinator groups on the phylogeny (Adams and Collyer 2018). We tested for evidence of phylogenetic signal in compound number, compound diversity, compound amount (total area of floral scent peaks), (*E*)-beta-ocimene production, and (*E*)-beta-caryophyllene production, detecting weak and non-significant phylogenetic signal overall (λ= 4.14795e-05, p=1). We also tested for evidence of phylogenetic signal in the multivariate floral scent dataset, detecting non-significant almost zero phylogenetic signal (Kmult=0.43, p=0.68). We therefore decided against using phylogenetic ANOVA with simulations based on Brownian motion and instead used a method of randomizing residuals in a permutation procedure (Adams and Collyer 2018). We found that compound number, compound diversity, compound amount, and (*E*)-beta-ocimene production were all significantly different between bee-pollinated and hummingbird-pollinated species (compound number, F_1,26_=7.2, p=0.008; compound diversity, F_1,26_=5.8, p=0.01; compound amount, F_1,26_=627.5, p=0.008; (*E*)-beta-ocimene production, F_1,26_=4.7, p=0.03). (*E*)-beta-caryophyllene production did not differ between pollinator groups (F_1,26_=1.0, p=0.4). Our comparative analyses complement our previous analyses, again showing that bee-pollinated species produce higher amounts of more diverse and complex floral scents and have higher (*E*)-beta-ocimene production.

To further investigate how floral scent has evolved across the *Costus* phylogeny, we carried out maximum likelihood ancestral state reconstruction of mean compound production. We overlaid the eight independent origins of hummingbird pollination represented by our dataset as determined in the full phylogenetic analysis of Vargas (2020) (Vargas et al. 2020). The ancestral state of all *Costus* species found in tropical America is to produce more compounds than average hummingbird-pollinated species (4.5 vs 2). In this reconstruction, the ancestral *Costus* population produced an average floral scent – more compounds than hummingbird-pollinated species but less than the average bee-pollinated species (Figure 3). Many hummingbird-pollinated species have since completely lost floral scent, as shown by the white circles (Figure 3). We see evidence for scent reduction or loss in all eight origins of hummingbird pollination in our dataset. In contrast, some bee-pollinated species, such as *Costus allenii*, have evolved much stronger floral scents than the ancestral state.

**Figure 3.**
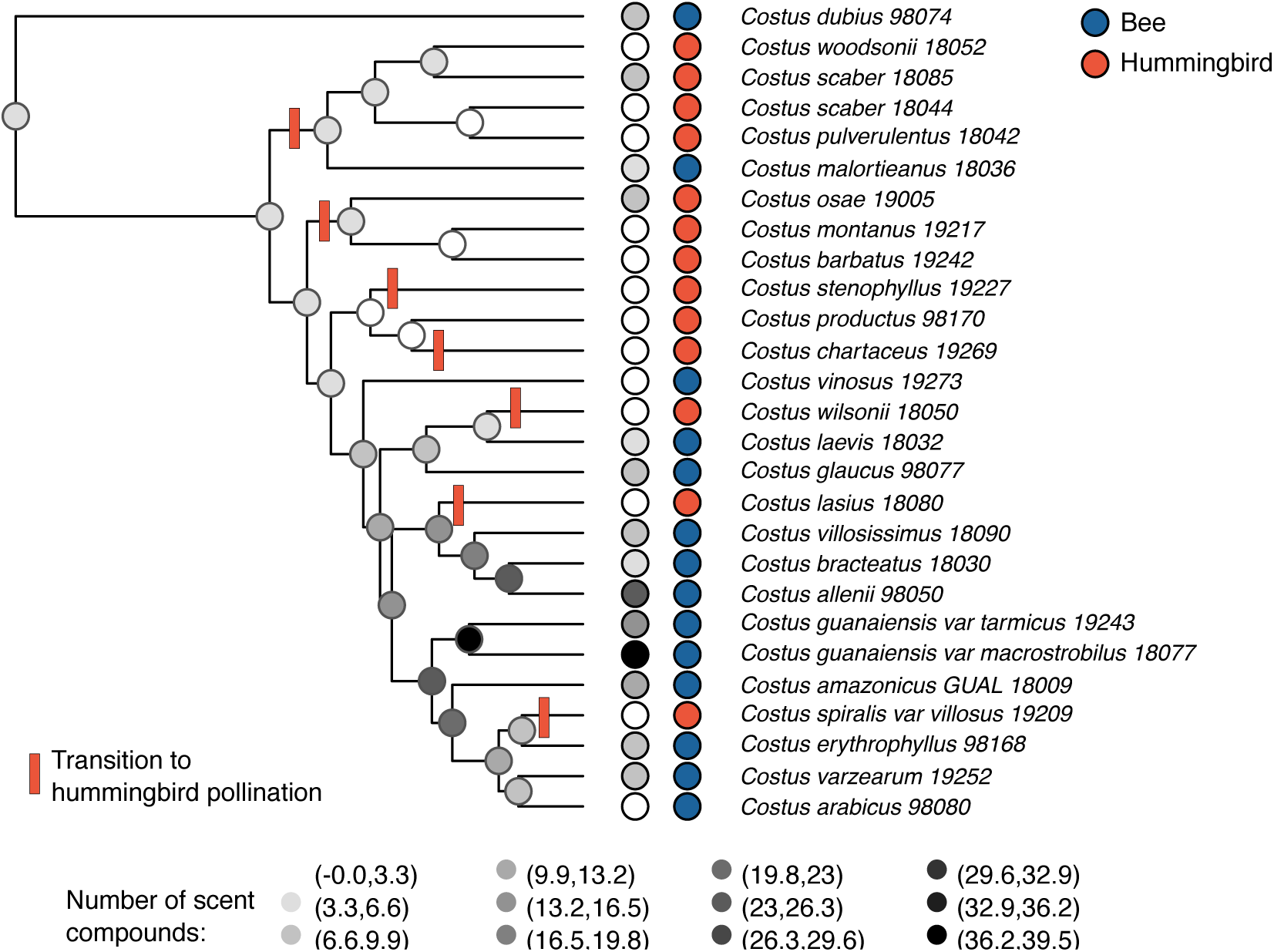
Ancestral state reconstruction of floral scent in the genus *Costus*. Pollinator for each species is illustrated by the right-hand dots. Mean compound number produced by each species is shown by the left-hand dots with each shade of gray representing a range of means. Darker dots indicate more compounds are produced. Predicted compound production at each node is shown following the same color scale. Tip labels following (Vargas et al. 2020). Transitions to hummingbird pollination were inferred from a larger species-level phylogeny (Vargas et al. 2020).

### Loss of floral scent is correlated with downregulation of floral terpene synthase genes

To analyze the genetic basis of floral scent evolution, we first chose a clade of closely related species with variation in floral scent production and pollination systems. We chose the clade containing *Costus lasius* due to the variation in floral scent production and the availability of a high-quality genome. In this clade, *Costus lasius* (hummingbird-pollinated) produces on average three floral scent compounds, whereas *Costus allenii* (bee-pollinated) produces 24 (one of the highest in our dataset), and *Costus villosissimus* (bee-pollinated) produces seven compounds (Figure 4A).

**Figure 4.**
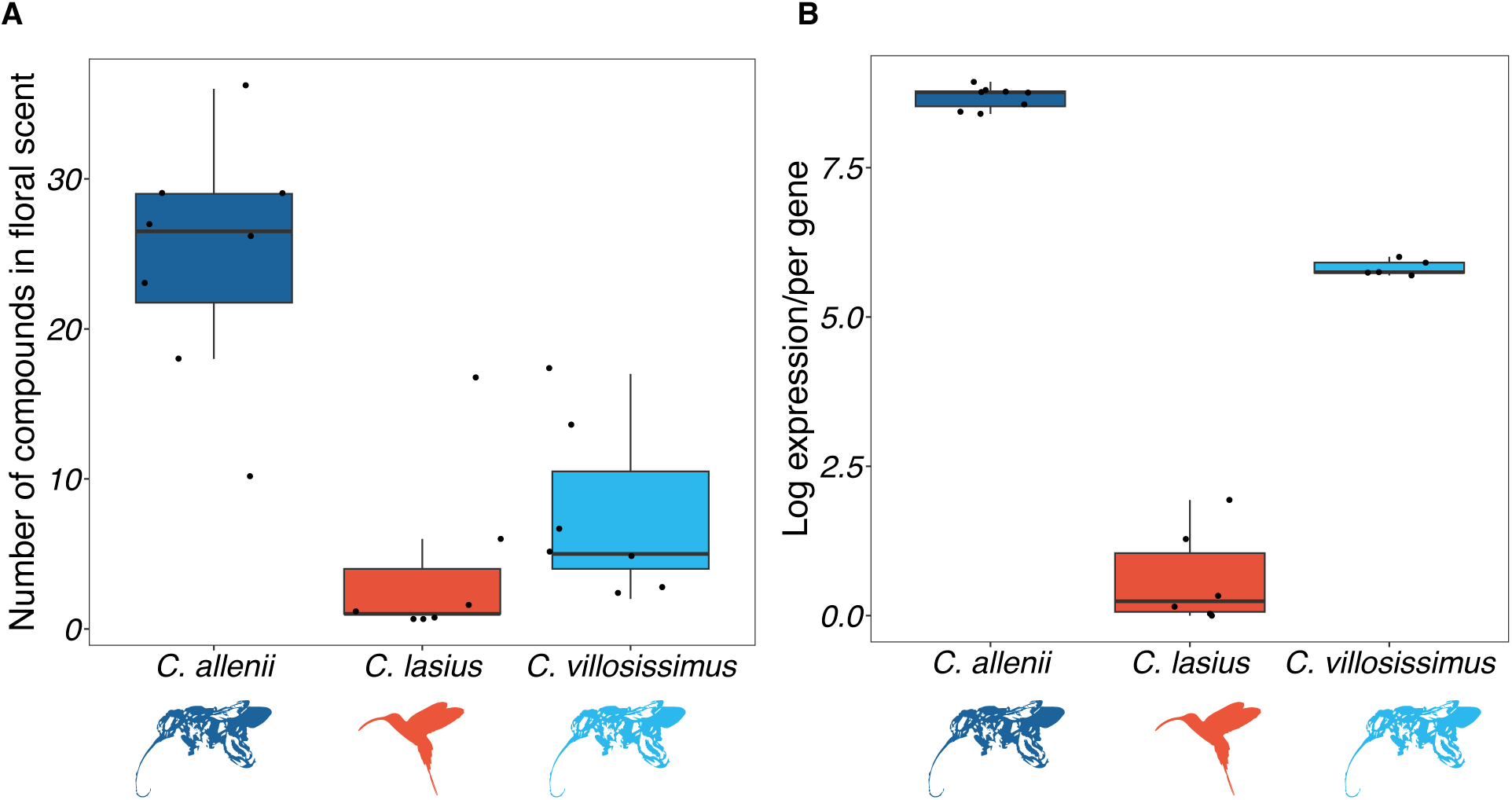
Differences in floral scent and gene expression between *C. allenii*, *C. lasius*, and *C. villosissimus*. Species differ significantly in A) number of floral scent components (ANOVA, F_2,19_=20.9, p<0.001), and (B) expression of terpene synthase transcripts (ANOVA, F_2,16_=508.4, p<0.001).

We first produced floral transcriptomes of all three species. After clustering transcripts with over 95% similarity, we found 112998 transcripts present in *C. allenii*, 80208 in *C. lasius*, and 96886 in *C. villosissimus*. Re-mapping rates were high in all three species: 99.2% in *C. allenii*, 99% in *C. lasius*, and 99.1% in *C. villosissimus*. All three also had BUSCO (Benchmarking Universal Single-Copy Orthologs) scores of more than 80% (Table S3), suggesting high-quality transcriptomes with most single-copy orthologs from the Embryophyta database present in all species.

Due to the dominance of terpenoid compounds in the floral scent, we focused on the expression of terpene synthase genes (TPS). TPS genes are responsible for producing terpenoid compounds. We found that 20 TPS genes were expressed in the floral transcriptome of *C. allenii*, compared to five in *C. villosissimus*, and only one in *C. lasius*, when expression was defined as one transcript per million (TPM) in at least one sample of the species. Furthermore, we found that counts of TPS expression per TPS gene expressed were much higher in both *C. allenii* and *C. villosissimus* when compared to *C. lasius* (Fig. 4B). In fact, in *C. allenii,* a TPS is the highest expressed gene in the floral transcriptome, and TPSs are also the 4^th^ and 6^th^ most expressed genes in the transcriptome, highlighting the extent of expression. On average, 12% of total expression can be attributed to TPSs in *C. allenii*, 0.17% in *C. villosissimus* and 0.00014% in *C. lasius*, highlighting the orders of magnitude differences between species.

### Floral scent loss is not driven by gene loss in *Costus*

While the transcriptome assemblies can give us insight into levels of TPS expression, it is hard to determine gene copy number due to the redundancies in the assemblies. For this, we need genome data to allow us to determine the number of TPS genes in each species. There are currently three publicly available *Costus* genomes: *C. bracteatus* (bee), *C. lasius (hummingbird)*, and *C. pulverulentus* (hummingbird) (Valderrama et al. 2022; Harenčár et al. 2023). *Costus lasius* and *C. pulverulentus* represent two independent transitions to hummingbird pollination (Kay and Grossenbacher 2022). To generate a dataset of two hummingbird- and two bee-pollinated species, we assembled an additional high-quality reference genome for bee-pollinated *C. allenii* (Table S4 for assembly statistics, Figure S3 to see alignment with *C. lasius* genome). We generated high-quality whole genome annotations for each species with high completeness and consistency (Table S5).

We annotated between 63-75% of each genome as repeat elements (Tables S6, S57, S8, S9). We found that, of the elements that could be classified, retrotransposable elements represented the largest percentage of the genome. These are the most abundant form of repeat elements found in angiosperm genomes. Many repeats were not classified into previously known families, perhaps suggesting the existence of new repeat elements.

We annotated a total of 66 TPSs in *C. allenii*, 44 in *C. bracteatus*, 40 in *C. lasius*, and 44 in *C. pulverulentus*. All TPSs belonged to previously described TPS families TPS-a, TPS-b, TPS-c, TPS-e/f, and TPS-g (Fig. 5, Fig. S4). Those TPSs responsible for floral scent biosynthesis are primarily found in the TPS-a, TPS-b, and TPS-g families. TPS-a members tend to produce monoterpenes, TPS-b members sesquiterpenes, and TPS-g members either monoterpenes or sesquiterpenes. We identified lineage-specific duplications in both TPS-a and TPS-b, as well as TPS-e/f in the bee-pollinated *C. allenii*. We did not find any evidence for pseudogenization or gene loss in the two hummingbird-pollinated species included in the gene family analyses, *C. lasius* and *C. pulverulentus*, with TPS genes containing complete protein domains and start codons.

**Figure 5.**
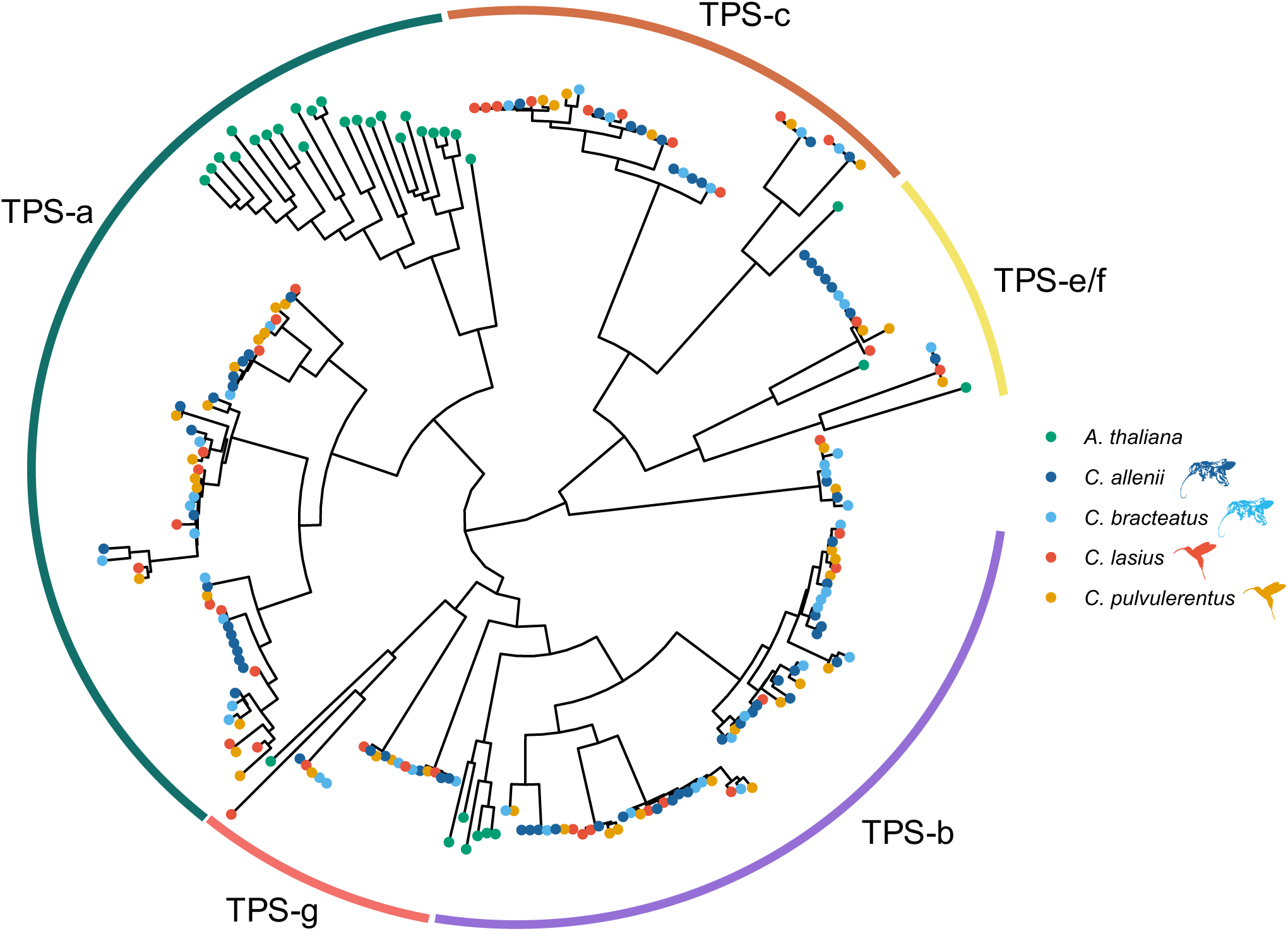
Phylogenetic analysis of terpene synthase gene families. We found the highest rates of gene gain and lowest rates of gene loss for *C. allenii* with the other three species exhibiting mixes of loss and gains across the gene tree. The phylogeny was constructed in IQ-TREE with 1000 bootstraps and model LG. TPS family classifications based on *Arabidopsis thaliana* are illustrated.

We used computational analysis of gene family evolution (CAFE5) to model gene gain and loss across the species phylogeny for each TPS clade identified in Figure 4 (Mendes et al. 2020). We found that *C. allenii* has gains in all five clades (TPS-a, TPS-b, TPS-c, TPS-e/f, TPs-g) with no evidence of losses, *C. bracteatus* shows no gains but losses in 4 clades, *C. lasius* and *C. pulvulerentus* both show gains in 1 and losses in 3. There is no clear correlation between pollination group and gene losses or gains with some evidence that *C. allenii* show enhanced gene duplication. We also estimated gene duplications and losses for each species using a gene tree-species tree reconciliation algorithm, reconcILS (Mishra et al. 2024). The algorithm reconciles the gene tree and species tree considering not only duplication and loss events but also incomplete lineage sorting (ILS). This is important for *Costus* due to the recent divergence times between species (Vargas et al. 2020). Using this approach, we identified the greatest number of duplication events for *C. allenii* (20), with 6 in *C. bracteatus*, and 4 in each hummingbird-pollinated species (*C. lasius* and *C. pulvulerentus*). We found the fewest loss events in *C. allenii* (2), with more losses in *C. pulvulerentus* (5), *C. bracteatus* (6), and *C. lasius* (11). We find support for increased lineage-specific duplications in *C. allenii*, however, our gene family analyses do not find evidence for increased gene losses in hummingbird-pollinated species.

We mapped the RNAseq reads both to the *C. allenii* genome and the *C. lasius* genome. In *C. allenii*, we found a consistent percentage of successfully mapped reads attributed to TPSs as with the transcriptome (12% transcriptome, 13% mapped to either genome). We found some differences for both *C. lasius* and *C. villosissimus*. For the *C. villosissimus* transcriptome mapping, we found 0.17% of reads were attributed to TPSs, in comparison to 0.009% when mapped to either genome. Due to a lack of genome for *C. villosissimus*, it is unclear whether this could be due to genetic changes or rearrangements not captured by the available genomes. For *C. lasius*, we found only 0.00014% of reads mapped to the transcriptome were due to TPSs, in contrast to 0.011% of the genome-mapped reads.

When *C. allenii* reads were mapped to the *C. allenii* genome, we found high levels of multi-mapping. This is due to recent TPS duplications in *C. allenii* which are highly expressed. To illustrate this, the highest gene expressed in *C. allenii* is *TPS49*, and the third most expressed “gene” consists of reads that multimap to *TPS49* and *TPS52*. This clade is represented by only one gene in each of the other species. Other recent lineage-specific duplications include *TPS2*, *TPS3*, *TPS4*, and *TPS5* where the majority of reads multimap due to sequence similarity between duplicates. Both of these clades have only one member in each of the other species with an available genome sequence (Fig S3). This highlights the complexity of mapping to gene families with recent duplications, especially when they are highly expressed, meaning that the typical choice of ignoring multi-mapped reads will lose large percentages of mapped reads from these duplications. We, therefore, decided to carry out our differential expression analysis using the *C. lasius* genome.

By mapping all reads to the *C. lasius* genome, we identified 4790 genes differentially expressed between *C. allenii* and *C. lasius*, 7678 between *C. allenii* and *C. villosissimus*, and 6191 between *C. lasius* and *C. villosissimus*. Of the 40 TPS genes found in the *C. lasius* genome, 15 were differentially expressed between *C. allenii* and *C. lasius*, with 13 of these upregulated in *C. allenii*. This includes *TPS33*, orthologous to the *TPS49*/*52* clade, the most highly expressed genes in *C. allenii*. We found 19 TPSs that were differentially expressed between *C. allenii* and *C. villosissimus*, with 14 upregulated in *C. allenii* and 5 in *C. villosissimus*. Ten TPS genes were differentially expressed between *C. lasius* and C*. villosissimus*, with 7 upregulated in *C. villosissimus* and 3 in *C. lasius*. Overall, more TPSs were upregulated in *C. allenii* when compared to either *C. lasius* or *C. villosissimus*. Our gene ontology analysis found little evidence for functional enrichment in differentially expressed genes between species (Table S10).

## Discussion

Repeated evolutionary transitions provide unique opportunities to study the extent of convergence both at the phenotypic and molecular level, shedding light on the predictability of evolution. Transitions to hummingbird pollination are widespread in the Americas and are predicted to be associated with loss of floral scent, however, this has not been studied in a comparative framework across multiple independent transitions within the same genus. In this study, we combine comparative chemical ecology with genomics and transcriptomics and demonstrate that floral scent in *Costus* is dominated by terpenes and its evolution is highly associated with pollinator type. We found that hummingbird-pollinated *Costus* species have lost or reduced their floral scent across eight independent evolutionary origins, whereas bee-pollinated species in the Americas exhibit either maintenance of floral scent or gain of a more diverse scent profile. We find that this is related to downregulation of genes associated with floral scent production (in this case terpene synthases) in floral tissues of hummingbird-pollinated species but is not accompanied by loss of terpene synthase genes.

Bee-pollinated *Costus* species showed much greater variation in their floral scents than hummingbird-pollinated species, with some having very diverse floral scents, yet others producing far fewer compounds. Bee-pollinated species also show greater morphological variation, possibly reflecting variation in orchid bee diversity (Kay and Grossenbacher 2022). Furthermore, not all species in our dataset follow the expected patterns. For example, *Costus arabicus* and *Costus vinosus* (bee-pollinated) produce little scent. It is unclear whether the variation in floral scent between different bee-pollinated species is related to pollinator preference, other abiotic or biotic factors, or historical contingency.

There are multiple hypotheses for why floral scent would be lost in hummingbird-pollinated species. Firstly, it could be because there is simply no selection to maintain the production due to poor sense of smell, or lack of preference shown for floral scent, by hummingbirds (Goldsmith and Goldsmith 1982; Knudsen et al. 2004; Driver and Balakrishnan 2021). If this were the case, we might expect to find a correlation between length of time following pollinator transition and extent of floral scent loss. However, our data shows that downregulation of floral scent can occur on relatively quick evolutionary timescales. The Latin American radiation of *Costus* is dated at 3 million years (95% CI, 1.50–4.87) and so all transitions to hummingbird-pollination have occurred within this timeframe (Vargas et al. 2020). *Costus wilsonii*, *C. lasius*, and *C. spiralis* are all hummingbird-pollinated species that are most closely related to bee-pollinated species, representing more recent evolutionary shifts to hummingbird pollination. Yet all three species produce little to no floral scent, suggesting there can be quick breakdown of floral scent production following pollinator transitions. This observation is consistent with previous studies that have shown that floral scent often evolves rapidly in response to selection (Ramos and Schiestl 2019; Zu et al. 2020; Liu et al. 2024).

The rapid and repeated loss of scent in hummingbird-pollinated species suggests a shared strong selection pressure against floral scent production. One potential mechanism is the energetic cost of terpene production. A similar argument has been made in the case of eye regression in cavefish. If vision is energetically costly to maintain, then it should be selected against when no longer useful (Moran et al. 2015; Sifuentes-Romero et al. 2023). This is supported by evidence of the energetic costs of maintaining a visual system (Moran et al. 2015). The metabolic cost of terpene production is unknown but is thought to be minimal (Raguso 2016; Pichersky and Raguso 2018). In fact, it has been argued that the complex network of terpene metabolism, in which individual enzymes produce multiple products, makes terpene pathways more efficient, compared with a linear pathway with separate enzymes and regulators for each step (Lanier et al. 2023).

An alternative scenario to the energetic cost is that terpene production has ecological costs in hummingbird-pollinated species. For example, there may be strong selection against bee attraction (Castellanos et al. 2004; Bergamo et al. 2016). Many benefits to hummingbird, compared with bee, pollination have been described. Birds do not consume pollen, leading to less pollen loss, and birds can travel longer distances between plants and promote higher gene flow, which is beneficial to combat inbreeding depression (Thomson and Wilson 2008; Krauss et al. 2017; Dellinger et al. 2022). One caveat is that comparisons in pollination effectiveness between bees and hummingbirds have not been carried out in orchid bees. Both male and female orchid bees tend to forage over long distances, perhaps resulting in a more “bird-like” pollen transport (Janzen 1971; Janzen 1981; Wikelski et al. 2010; Opedal et al. 2017; Kay and Grossenbacher 2022; Gamba and Muchhala 2023). While pollinator efficiency is unknown in *Costus*, rapid floral scent loss is congruent with the hypothesis of “bee-avoidance” following establishment of hummingbird pollination.

A final piece of evidence to support bee-repellence is the large reduction in beta-ocimene production in hummingbird-pollinated flowers relative to bee-pollinated flowers, as this compound is a common insect pollinator attractant (Farré-Armengol et al. 2017). In *Mimulus*, loss of monoterpenes, including beta-ocimene, maintains reproductive isolation between bee- and hummingbird-pollinated species (Byers et al. 2014). Interestingly, we found that α- and β-pinene were produced by both bee- and hummingbird-pollinated species. One potential reason is that pinenes do not repel orchid bees but do repel other types of bees. Terpenoid-dominated scents have been found to repel honey bees (Larue et al. 2016), and in particular α-pinene has a repellent effect on honey bees (Fernandes et al. 2019). Furthermore, α-pinene has been found in other hummingbird-pollinated species with a suggested role as a moth pollinator deterrent (Bischoff et al. 2014). We speculate that bee-pollinated species emit a mix of compounds, some of which potentially attract orchid bees, and some of which potentially repel other pollinators to narrow attraction to orchid bees, specifically. Then when floral scent is reduced in hummingbird-pollinated flowers, the compounds which are attractive to orchid bees are lost but not those which are general bee repellants. Further experimental work is needed to test these hypotheses.

We found that the floral scents of *Costus* are dominated by terpene compounds. Although few Zingiberales have described floral scents, our results are consistent with previous reports from the genus *Hedychium* that also found terpene-dominated floral scents (Báez et al. 2011; Zhou et al. 2022). Terpene diversity in a given organism is mainly determined by the terpene synthase (TPS) gene family, which carries out crucial steps in terpene formation (Tholl 2006). We did not find evidence that hummingbird-pollinated species with an available genome sequence (*C. lasius* and C. *pulverulentus*) exhibited increased rates of TPS gene loss than the bee-pollinated species. There are multiple reasons why trait loss would be associated with changes in gene regulation rather than gene loss. While *C. lasius* represents a more recent transition to hummingbird pollination than *C. pulvulerentus*, both transitions have occurred within the last ∼3 million years (95% CI, 1.50–4.87)as *Costus* radiated in Latin America (Vargas et al. 2020). There may not have been enough time for pseudogenization to occur, a process estimated to take 0.5-6 million years (Marshall et al. 1994; Esfeld et al. 2018). Potentially, the genes will eventually be lost but are currently in the first phase, reduction in expression, which according to the “step-wise” model to pseudogenization is then followed by relaxed selection and then loss-of-function mutations (Graham et al. 2023).

Alternatively, we might not expect any TPS genes to be lost even in the long term. Terpenes, and therefore TPSs, are important for many other roles in plants. Plants use terpenes to deter herbivores, to attract parasitoids or predators of herbivores, and as anti-fungal and anti-bacterial compounds to prevent disease (Gershenzon and Dudareva 2007). These pleiotropic functions of TPSs could explain why we observe downregulation in floral tissues specifically rather than any pseudogenization. One argument against this is that in other systems TPS pseudogenization does occur following pollinator transitions. In *Mimulus*, loss of monoterpene production (specifically beta-ocimene), is not due to expression differences but loss-of-function mutations (Peng et al. 2017). It is unclear whether pseudogenization will occur at some point in *Costus*.

Although we did not observe gene loss in hummingbird-pollinated lineages, we did observe extensive lineage-specific duplications in the bee-pollinated species *C. allenii*. *Costus allenii* was one of the species with highest scent diversity in our dataset, and we found multiple TPS genes to exhibit several recent lineage-specific duplication events. We also found extremely high TPS expression, accounting for approximately 12% of expression in floral tissue. The gene with the highest expression in floral tissue was found to be a TPS-a family member, with high expression also found also for several TPS-b members. These two families produce sesquiterpenes and monoterpenes, respectively, both of which are found in the floral scent of *C. allenii*. In *C. lasius* and *C. villosissimus*, we found reduced TPS expression compared to *C. allenii*, as expected based off their reduced floral scent production. We also saw high levels of phenotypic variation in *C. villosissimus*, and there is no publicly available genome, making interpretation of the results more difficult. We speculate that *C. allenii* has higher floral scent production than closely related *C. villosissimus* because *C. allenii* grows as isolated individuals in deeply shaded, perennially wet forest, whereas *C. villosissimus* grows at forest edges and large treefall gaps, and often has conspecifics nearby (Chen and Schemske 2015). Thus *C. allenii* may need more scent to attract pollinators. While the overall trends are clear, we hesitate to over-interpret the gene expression data as we have only sampled at one timepoint and it is possible that there is floral scent production happening earlier that is not captured in our data.

We observe more consistency and predictability in the phenotypic patterns than the underlying molecular mechanisms. In many systems, parallel changes in floral scent have accompanied independent pollinator shifts, including the current study (Knudsen et al. 2004; Byers et al. 2014). This can also be seen in other reproductive shifts, such as transitions to self-fertilization (Wozniak et al. 2022). In some systems, the underlying molecular mechanisms of floral scent loss are also convergent, having been shown to involve the same loci in multiple independent evolutionary events. For example, loss of floral scent in different *Mimulus* lineages is due to changes at the same loci, including frameshifts, deletions, and potentially post-transcriptional regulation (Peng et al. 2017). A similar pattern was found in two different systems that have lost benzaldehyde floral scent production due to either evolution of self-fertilization (Sas et al. 2016), or transition to hummingbird pollination (Amrad et al. 2016). The same locus contains loss-of-function mutations in both cases (Raguso 2016). However, this is not universally true. In *Capsella*, the loss of beta-ocimene following transition to self-fertilization in one lineage is due to changes in subcellular enzyme localization, but QTL results suggests another locus is responsible for scent loss in a related species (Wozniak et al. 2022). In this study, we saw no evidence for pseudogenization. Instead, we found evidence that gene expression differences are responsible for scent differences between *Costus* species, with scent loss likely due to changes in, potentially tissue-specific, gene expression.

In summary, we have shown that *Costus* species exhibit repeated loss of floral scent following transitions from bee to hummingbird pollination, suggesting a shared strong selection pressure favoring scent loss. We also find bee-pollinated species that have recently gained more diverse scents, which, at least in *C. allenii*, is associated with lineage-specific terpene synthase duplications. We show that scent loss is not associated with loss of terpene synthase genes but rather appears to be due to gene downregulation. This demonstrates the capacity for rapid metabolic changes in response to selection following pollinator transitions. This system provides us with the exciting opportunity to identify the cis- or trans-regulatory changes driving gene expression differences in multiple independent transitions to hummingbird pollination, allowing us to study the extent of genetic convergence.

## Materials and Methods

### Floral scent sample collection

We collected 100 samples of 30 species of *Costus* between 2019 and 2023 (Table S1). We sampled an average of 4 individuals per species for bee-pollinated species, and 3 for hummingbird-pollinated species. We, therefore, expect that missing compounds due to under-sampling will occur to a similar extent for both bee- and hummingbird-pollinated species. All samples for *Costus glaucus*, *Costus laevis*, *Costus montanus*, *Costus stenophyllus*, and *Costus wilsonii* were collected in Costa Rica (La Gamba, Las Cruces, Las Alturas, and Monteverde). All other species were sampled in the UCSC greenhouses in Santa Cruz, California, USA from plants composing the UCSC living *Costus* collection. For each day of sampling, both floral scent samples and ambient samples were collected (with one exception: a *Costus scaber* sample collected on April 8^th^ 2019 was missing a control and so we used a control from the nearest date available, March 7th). Scent was sampled from a single flower using dynamic headspace sampling from approximately 9 am until 3 pm (6 hours). This time point was chosen as it is the window of most pollinator activity during the lifespan of the flower (Kay and Schemske 2003). Flowers from living plants were placed inside oven bags (Reynold’s). A Spectrex PAS-500 air pump with Tygon tubing was used to pull scented-air through a volatile trap for 6 hours at a flow rate of 200 ml air / min. The volatile trap contained a filter of glass wool and 50 mg Porapak. We conditioned scent traps by passing 5 mL of hexane before usage. Volatiles were eluted from the trap using 400μl of hexane and stored at -20°C.

### Chemical analysis

Samples were analyzed using Agilent model 5977A mass-selective detector with an Agilent GC model 7890B. Samples were run on a HP-5 Ultra Inert column (Agilent, 30m x 0.25mm, 0.25 μm). For analysis, one microliter of each sample was injected in splitless mode using the ALS 7694 autosampler with helium as the carrier gas (250°C injector temperature). The program started at 40 °C for 3 minutes and then rose at 5 °C/min to 210°C. The temperature was held at 210°C for 1 minute, before rising at 20°C/min to 300°C where it was held for 2.5 minutes, and 315°C for a final minute.

Compounds were identified by comparing mass spectra and retention indices with reference libraries. The identity of some compounds was confirmed by comparison to authentic reference standards (see Table S2). Compounds that were not found in at least two samples were removed. A two-step process was used to filter datasets for floral scent components. First, compounds were filtered to include only those which are found in five times higher amounts in floral than ambient samples using the “filter_ambient_ratio” function from the *bouquet* package (Powers et al. 2023). This resulted in a list of ambient compounds which were not considered floral scent compounds. Second, using the raw data matrices, we subtracted the ambient compound amounts from the corresponding floral scent amount collected on the same date. We then removed the ambient compounds identified by the package *bouquet* from this subtracted matrix to create the final data matrix of floral scent.

### Statistical analyses of floral scent

To visualize divergence in floral scent among samples we used nonmetric multidimensional scaling (NMDS) using the “metaMDS” function in *vegan* (Oksanen et al. 2020). To test for differences both between pollination groups, and between floral and ambient samples, we carried out a PERMANOVA (permutational multivariate analysis of variance) using the “adonis2” function in *vegan* (Bray–Curtis distance matrix, 1000 permutations). To identify which groups were significantly different, we carried out post hoc pairwise testing using the “pairwise.perm.MANOVA” function in the *RVAideMemoire* package with a Bonferroni correction (Hervé 2021). We did this for both filtered and unfiltered datasets.

We tested for differences in compound number produced by bee-pollinated and hummingbird-pollinated species using ANOVA with a nested model including species within pollinator group. To test for differences in the diversity of compounds produced by bee-pollinated and hummingbird-pollinated species, we used the package *chemodiv* (Petrén et al. 2023). This package considers the biochemical and structural diversity of the compounds present in a sample to calculate the overall chemodiversity per sample. First, we used the “calcDiv” function to calculate the functional Hill diversity index for each sample. Then we carried out an ANOVA test for the diversity indices, using a nested model of species nested within pollinator group to test for differences in chemodiversity. We also tested for differences in compound production using ANOVA. In this case a proxy for compound amount was used: the sum of the total area of floral scent peaks in the sample, or the total ion abundance in the sample. To test for difference in compound frequency between bee- and hummingbird-pollinated samples, we carried out chi-squared tests and adjusted the p-values using the “p.adjust” function in R.

### Phylogenetic analyses of floral scent

To investigate the evolution of floral scent in a phylogenetic context we used a previously published phylogeny (Vargas et al. 2020). First, we trimmed the phylogeny to include only those species for which we have scent data using the “drop.tip” function in *ape* (Paradis and Schliep 2019). To test for correlation between pollination group and the phylogenetic covariance matrix, we used a two-block partial least squares analysis using the “two.b.pls” function in the *geomorph* package (Adams and Otárola-Castillo 2013; Adams and Collyer 2018). A significant result means that our power to test for differences between groups using phylogenetic simulation-based is weakened due to group aggregation on the phylogeny. We then used the “phylosig” function in the package *phytools* to test for phylogenetic signal in multiple traits (compound number, compound diversity, compound amount, (*E*)-beta-ocimene production, and (E)-beta-caryophyllene production) (Revell 2012). We also tested if floral scent data overall shows phylogenetic signal using the multivariate generalization of Bloomberg’s K (K_mult_) (Adams 2014). Under Brownian motion, K_mult_ is expected to be 1. If K_mult_>1, higher phylogenetic signal is detected than the null expectation under Brownian motion. If K_mult_<1, lower phylogenetic signal is detected than the null expectation under Brownian motion. To calculate the K_mult_ of *Costus* floral scent, we used the filtered dataset with 1000 iterations to test significance. We carried out the analysis using the “physignal” function in the *geomorph* package (Adams and Otárola-Castillo 2013). Due to the weak phylogenetic signal detected in both multivariate and univariate analyses, we used the “procD.pgls” function in the *RRPP* package to test for differences in traits between pollinator groups rather than a simulation-based ANOVA (Adams and Collyer 2018; Collyer and Adams 2018). This function uses a method of randomizing residuals in a permutation procedure to evaluate the significance of phenotypic differences between groups considering the phylogeny and has higher statistical power than methods using phylogenetic simulations (Adams and Collyer 2018). We set lambda at 4.14795e-05 as calculated for the univariate variables. We corrected for multiple testing using the “p.adjust” function in R.

To better understand how floral scent has evolved in the genus *Costus*, we carried out an ancestral state reconstruction of the number of compounds produced. The species-level phylogeny from (Vargas et al. 2020) contains 24 hummingbird-pollinated species, 25 bee-pollinated species, and 3 bee-pollinated African outgroups. Our subsampled phylogeny contains samples in proportion to the full phylogeny (13 hummingbird-pollinated species, 13 bee-pollinated species, and 1 bee-pollinated African outgroup). We note that there are limitations to this sub-sampled phylogeny but we find value in using it to infer general trends and not values at specific nodes. To do this, we used the function “ace” in the *ape* package for ancestral character estimation (Paradis and Schliep 2019). We divided the compound number trait into twelve categories, each of which were assigned a different shade of blue. We plotted both the compound number for each phylogenetic tip label and internal node to illustrate how floral scent has evolved in *Costus*. We overlaid the origins of hummingbird pollination as described in (Vargas et al. 2020).

### Sample collection for RNAseq

Live individuals of *Costus allenii* and *Costus villosissimus* were collected in Soberanía National Park, Colón Province, Panama. Vouchers are deposited at University of Panama (*Costus allenii*: 0132264, 0132265; *Costus villosissimus*: 0132270). Live individuals of *Costus lasiu*s were collected in El Valle de Antón, Cocle Province, Panama. Voucher is deposited in Michigan State University (MSC) herbarium (Kay 0321). For all three species, the individuals used for RNAseq analyses were grown in greenhouses at UCSC. Freshly opened flowers of each species were removed at approximately 9 am and flash frozen immediately in liquid nitrogen before being stored at -80°C until extraction. Flowers of *Costus* open early in the morning and fall off by mid-afternoon.

### RNA extraction and sequencing

Frozen flowers were removed and dissected on dry ice. Tissue from the petals and labella was broken down using a TissueLyser (Qiagen). RNA was then extracted using an RNeasy Mini Kit (Qiagen) with Qiashredder spin columns (Qiagen) following standard protocols. Samples were sent to Novogene for quality control, library preparation, and sequencing. Our final dataset consisted of 19 samples: 8 *Costus allenii* (from 4 individual plants), 6 *Costus lasius* (from 2 individual plants), and 5 *Costus villosissimus* (from 1 individual plant). Samples were sequenced on a NovaSeq 6000 using 150 bp paired end reads. This generated approximately 53 million reads per library (mean=52.74 million, SD=12.01 million, N=19). The raw sequence data were deposited in the SRA under the BioProject accession PRJNA1010186.

### Transcriptome assembly

Before assembling transcriptomes, we trimmed the raw reads using TrimGalore! (Martin 2011). We then assembled transcriptomes for each species using Trinity version 2.5.1 with default settings (Grabherr et al. 2011; Haas et al. 2013). To reduce redundancy, we clustered contigs with a minimum sequence identity of 95% using CD-HIT version 4.6.8 (Fu et al. 2012). We assessed transcriptome completeness and quality using BUSCO (v5.6.1) (embryophyta_odb10 dataset) (Simão et al. 2015; Manni et al. 2021; Zdobnov et al. 2021). We also checked assembly quality by mapping RNAseq reads back to the assembly using Bowtie (v2.4.1) to calculate mapping rates (Langmead and Salzberg 2012).

### Terpene synthase gene expression analysis

We extracted the longest isoforms for each Trinity “gene” using the Trinity script “get_longest_isoform_seq_per_trinity_gene.pl” and translated these sequences. We then searched for potential TPSs using hmmsearch two HMM profiles: Terpene_syth_C (PF03936) and Terpene Synthase N-terminal domain (PF01397). We used the R package rhmmer to parse the output from hmmsearch resulting in a list of putative TPS sequences for each species. To reduce the TPSs to only those expressed in the dataset, we used a threshold of TPM>1 (transcripts per million) in at least one individual of a species (Eddy 2011; Arendsee 2017). To calculate expression of each putative TPS we normalized counts using Gene length corrected trimmed mean of M-values (GeTMM), an approach that improves inter- and intra-sample comparisons (Smid et al. 2018). To do this, we first calculated FPKM (fragments per kilobase of transcript per million fragments mapped) for each transcript and carried out TMM-normalization using the package *edgeR* (Robinson et al. 2010). We then combined GeTMM values for transcripts which were identified by Trinity to be the same gene. This does not normalize counts between species but is used to determine whether genes are lowly or highly expressed within a species.

### *Costus allenii* genome assembly

We selected a wild-collected individual *C. allenii* from Pipeline Road, Soberanía National Park, Colón Province, Panama for *de novo* genome assembly. A voucher for this population of *C. allenii* is deposited at the University of Panama Herbarium (PMA:0132264). This individual was grown to flowering in the UCSC greenhouses from a cutting, and we harvested 1.5 g fresh tissue from leaf meristems and flower buds. We isolated cell nuclei following Workman (Workman et al. 2019) and then used the Circulomics Nanobind Plant Nuclei Big DNA kit to extract HMW DNA. We assessed DNA concentration with a Qubit Fluorometer (Thermo Fisher Scientific), purity with 260/280 and 260/230 absorbance ratios from a NanoDrop Spectrophotometer (Thermo Fisher Scientific) and ran a 7% agarose gel at 70 V for over 8 h to assess DNA fragment integrity and size. HMW DNA was sent to the UC Davis Genomics Center for additional cleaning, Pacific Biosciences HiFi library preparation, and sequencing on one PacBio Revio SMRT cell. This produced 74.8 Gb of HiFi yield and 7.9 M HiFi reads. We assembled the PacBio data into a primary and alternative assembly using HiFiasm (v0.19.8)(Cheng et al. 2021). We then used RagTag (v2.1.0)(Alonge et al. 2022) to scaffold *C. allenii* contigs using the closely related *C. lasius* as a reference. We used D-GENIES (Cabanettes and Klopp 2018) with Minimap2 (Li 2018) to generate a dotplot comparing the two assemblies both before and after scaffolding. The genome assembly quality was assessed using BUSCO v 5.6.1 (Simão et al. 2015). Raw sequence data and the final genome assembly were submitted to NCBI under BioProject PRJNA1091680.

### Genome annotations

Of the three species for which we generated RNAseq data, an unannotated genome was previously available for *C. lasius* (Harenčár et al. 2023). In this study we also generated a genome assembly for *C.allenii*. In addition, previously published genomes are available for both *C. bracteatus* and the more distantly-related *C. pulverulentus* (Valderrama et al. 2022; Harenčár et al. 2023). We started by processing RNAseq data as evidence for annotations. Trimmed reads from *C. lasius* were mapped using 2pass mapping in STAR to the *C. lasius* genome, and the *C. pulverulentus* genomes (Martin 2011; Dobin et al. 2013). Bam files were concatenated for use as described below. Trimmed reads from *C. allenii* were mapped using 2pass mapping in STAR to both the *C. allenii* and the closely related *C. bracteatus* genomes, and bam files concatenated per species (Martin 2011; Dobin et al. 2013).

We annotated the genomes using BRAKER3 (v3.0.7) which combines RNA-seq and protein data in an automated pipeline (Lomsadze et al. 2005, 2014; Stanke et al. 2006, 2008; Gotoh 2008; Iwata and Gotoh 2012; Buchfink et al. 2015; Hoff et al. 2016, 2019; Kovaka et al. 2019; Brůna et al. 2020, 2021; Pertea and Pertea 2020; Gabriel et al. 2023). This pipeline requires a masked genome. We identified and annotated tandem repeats and transposable elements using RepeatModeler (Flynn et al. 2020) (v.2.0.4 for *C. lasius* and v.2.0.5 for the other genomes) and RepeatMasker v.4.1.5 (Tarailo-Graovac and Chen 2009). We then used RepeatMasker with the outputs of RepeatModeler to generate softmasked genomes. The masked genomes were then provided to BRAKER along with plant proteins downloaded from OrthoDB (odb10_plants) (Zdobnov et al. 2021). For all species, BRAKER was run in ETP mode which combines evidence from both proteins and RNAseq (Gabriel et al. 2021). BUSCO v 5.6.1 (Simão et al. 2015) and OMArk (Nevers et al. 2022) were used to evaluate the completeness and consistency of the sets of gene models. During the project, the *C. lasius* genome was updated (direction of arms on chromosome 1 were altered) and so we re-scaffolded chromosome 1 of *C. allenii* and transferred our annotation using liftoff (Shumate and Salzberg 2021).

### Terpene synthase gene family annotation

Following whole-genome annotation, we manually curated the gene models for terpene synthase (TPS) genes in the annotation. To do this, we used bitacora (Vizueta et al. 2020), a pipeline which curates an existing annotation and finds additional gene family members. In this pipeline, BLASTP and hmmer are used to search for gene family members of interest within the existing annotation (Altschul et al. 1990; Eddy 2011). Additional regions of the genome are then searched using TBLASTN and are annotated using GeMoMa (Keilwagen et al. 2019). Hmmer is used to validate the gene models. As input, we used two HMM profiles: Terpene_syth_C (PF03936) and Terpene Synthase N-terminal domain (PF01397) downloaded from Pfam. In addition, we provided previously annotated genes from *Arabidopsis thaliana* and *Oryza sativa* (Chen et al. 2011; Yu et al. 2020; Jia et al. 2022). Gene models from bitacora and the genome-wide annotation from BRAKER were compared and TPS genes were manually curated using IGV (Robinson et al. 2011). We also searched for additional TPSs by carrying out exonerate searches using the curated *Costus* gene models (Slater and Birney 2005). We checked all annotated TPSs for complete protein domains using the NCBI conserved domain search (Marchler-Bauer et al. 2015). We excluded proteins with less than 250 amino acids.

### Gene family analysis

To classify these TPSs in previously described TPS families, we constructed a phylogenetic tree based on amino acid sequences with TPSs from *Arabidopsis thaliana*. The *Costus* TPS protein sequences, combined with those from *A. thaliana*, were aligned with MAFFT v7.508 (-maxiterate 1000, using L-INS-I algorithm) (Katoh and Standley 2013). We then trimmed the alignment using trimAl (-gt0.6, sites only included when present in 60% of sequences) (Capella-Gutierrez et al. 2009). We used this alignment to construct a gene tree with IQ-TREE and the ModelFinder function to determine the best-fit model (Nguyen et al. 2015; Kalyaanamoorthy et al. 2017; Hoang et al. 2018). We rooted the tree using the *phytools* package in R (“midpoint.root” function)(Revell 2012; R Core Team 2023). We plotted the resulting gene trees using the following packages in R: *ape* (Paradis and Schliep 2019), *evobiR* (Blackmon and Adams 2015), *ggnewscale* (Campitelli 2023), *ggstar* (Xu 2022), *ggtree* (Yu et al. 2017), *ggtreeExtra* (Xu et al. 2021), and *tidytree* (Yu 2022).

To investigate patterns of gene family evolution we used computational analysis of gene family evolution CAFE (v5.1) which uses a birth-death model of gene family evolution to model gene gain and loss across a species tree (Mendes et al. 2020). The required input files are a species tree which we adapted from (Vargas et al. 2020), and gene counts for each species in each clade (as identified in Fig. 5).

We also took an alternative approach by using reconcILS to reconcile our gene tree and species tree to estimate gene duplications and losses for each species (Mishra et al. 2024). The algorithm considers not only duplication and loss events but also incomplete lineage sorting (ILS), particularly relevant in this case due to the recent divergence times between *Costus* species (Vargas et al. 2020). To create a gene tree, we aligned all *Costus* TPS nucleotide sequences with MAFFT v7.508 (-maxiterate 1000, using L-INS-I algorithm) (Katoh and Standley 2013). The alignment was trimmed using trimAl (-gt0.6, sites only included when present in 60% of sequences) (Capella-Gutierrez et al. 2009) and used to construct a gene tree with IQ-TREE and the ModelFinder function to determine the best-fit model (Nguyen et al. 2015; Kalyaanamoorthy et al. 2017; Hoang et al. 2018). We rooted the tree using the *phytools* package in R (“midpoint.root” function)(Revell 2012; R Core Team 2023). The species tree was adapted from (Vargas et al. 2020). We used the default parameter settings.

### Differential expression analyses

First, we trimmed the reads using TrimGalore! (Martin 2011). We then mapped the reads from all three species to both the *Costus allenii* and *Costus lasius* genome (Harenčár et al. 2023). We ran 2pass mapping using STAR (Dobin et al. 2013). In general, we found higher levels of multi-mapped *C. allenii* reads when mapped to *C. allenii*. To investigate this we used mmquant (with merge, -m, set to 10) to count the reads, a tool which can account for multi-mapping reads (Zytnicki 2017). The output from mmquant was read into R for further analysis. To calculate percentage of gene expression due to TPS expression, we normalized reads by library size using TMM (trimmed mean of M values) (Robinson and Oshlack 2010). Due to the consistency in TPS expression as calculated by mapping to both genomes, we decided to use the *C. lasius* genome assembly to identify genes differentially expressed between the three species to remove the complication of multi-mapping. To do this, we first counted reads using featureCounts (Liao et al. 2014) and again read the output into R. We filtered genes to include those with at least one count per million (CPM) in at least one library and normalized reads by library size using TMM (trimmed mean of M values). Differential gene expression was evaluated using the non-parametric “noiseqbio” function from the *NOISeq* package (Tarazona et al. 2011; Tarazona et al. 2015). We used the default cut-off of q=0.95 in the “degenes” function which is equivalent to an adjusted p-value of 0.05.

### GO Analysis

We functionally annotated *C. alennii* genes using Blast2GO (Götz et al. 2008; Burge et al. 2012). We imported these functional annotations into R to carry out enrichment analysis with the package *topGO* (Alexa and Rahnenfuhrer 2022). We considered the background to be all genes included in that specific differential expression analysis after filtering.

### Plotting and data manipulation

Additional packages used for general plotting: *cowplot* (Wilke 2020), *ggplot2* (Wickham 2009), and *scico* (Pedersen and Crameri 2023). Packages used for data transformation and manipulation include: *data.table* (Barrett et al. 2024), *dplyr* (Wickham et al. 2021), *knitr* (Xie 2023), *magrittr* (Bache and Wickham 2023), *readr* (Wickham et al. 2023), *reshape2* (Wickham 2007), and *tibble* (Müller and Wickham 2022). Analyses were carried out in R version 4.3.2 (R Core Team 2023).

## Data availability

Data and R scripts used for analysis are available from Open Science Framework: https://osf.io/2ap4k/?view_only=6b8f5ff7d2ae499b871e1b1fb6fe6219. The raw sequence reads and final genome assembly for *Costus allenii* are deposited in the SRA (BioProject PRJNA1091680) https://dataview.ncbi.nlm.nih.gov/object/PRJNA1091680?reviewer=p4jo1ioj3vqualqk1kvege7aq l. The raw RNAseq data is deposited in the SRA (BioProject PRJNA1010186) https://dataview.ncbi.nlm.nih.gov/object/PRJNA1010186?reviewer=fil9vpq6up5g3pj7e654rtokje

## Acknowledgements

This work was funded by the National Science Foundation Dimensions of Biodiversity grant awarded to K.M.K. (DEB-1737889) and S.R.R. (DEB-1737771), and the Jean H. Langenheim Chair in Plant Ecology and Evolution held by K.M.K. We thank the Organization for Tropical Studies, Estación Biológica Monteverde, and Estación Tropical La Gamba for facilitating field work in Costa Rica and the Smithsonian Tropical Research Institute for facilitating field work in Panama. All field research was conducted with appropriate research permits in Costa Rica (M-P-SINAC-PNI-ACAT-026-2018, ACC-PI-0272018, MPC-SINAC-PNI-ACLAP-020-2018, INV-ACOSA-076-18, MPC-SINAC-PNI-ACTo-020-18, CONAGEBIO R-056-2019OT and R-058-2019-OT) and Panama (SE/AP-13-19, 09-99) We thank Julia Harenčár for assistance with lab work, Pedro Juarez and Cecilia Girvin for assistance with field sampling, and Alyssa Kaatmann, Aubrie Tait, Kate Uckele, Melina Sauerman Carrizosa, and Selena Vengco for assistance with greenhouse sampling. We thank J. Velzy and S. Childress for greenhouse care of the UCSC living *Costus* collection. We thank the Ramírez lab for helpful discussions.

## Author Contributions

Conceptualization, K.D., K.M.K., and S.R.R.; Investigation, K.D. and K.M.K.; Formal Analysis, K.D.; Visualization, K.D.; Resources, K.M.K. and S.R.R.; Writing – Original Draft, K.D.; Writing – Review and Editing, K.D., K.M.K., and S.R.R.; Supervision, K.M.K. and S.R.R.; Project Administration, K.M.K. and S.R.R.; Funding Acquisition, K.M.K. and S.R.R.

