## Supplementary material for "The convergent evolution of hummingbird pollination results in repeated floral scent loss through gene downregulation"

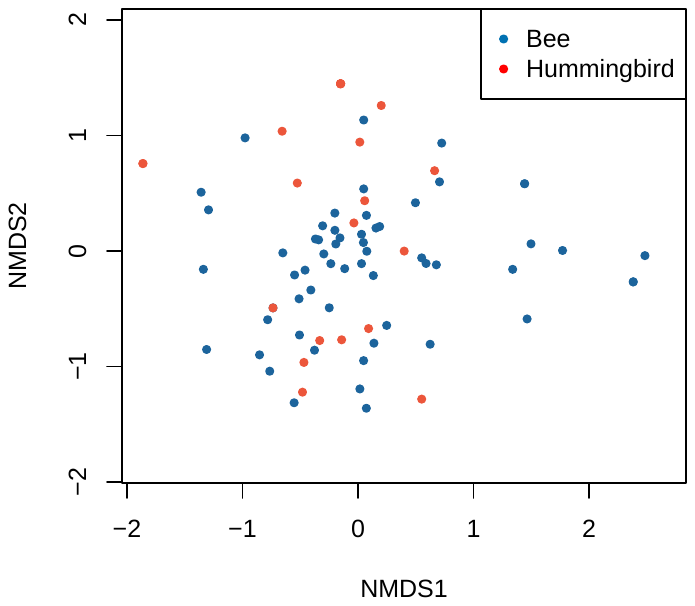

Figure S1. NMDS (nonmetric multidimensional scaling) plot illustrating the variation in the final filtered dataset (Stress=0.13) (PERMANOVA, pollination group: F_1,80_=4.2, p<0.001; species: F_26,80_=2.4, p<0.001; Pairwise PERMANOVA bee- vs hummingbird-pollinated species p=0.001).

Figure S2. Differences in number of scent compounds produced for bee-pollinated and hummingbird-pollinated *Costus* species collected in A) the greenhouse (ANOVA, pollination group: F_1,52_=38.7, p<0.0001, species: F_23,52_=8.1, p<0.0001) and B) the field (ANOVA, pollination group: F_1,18_=21.4, p<0.001, species: F_3,18_=2.2, p=NS). The species included in each are non-overlapping, which could explain some of the difference in scale.

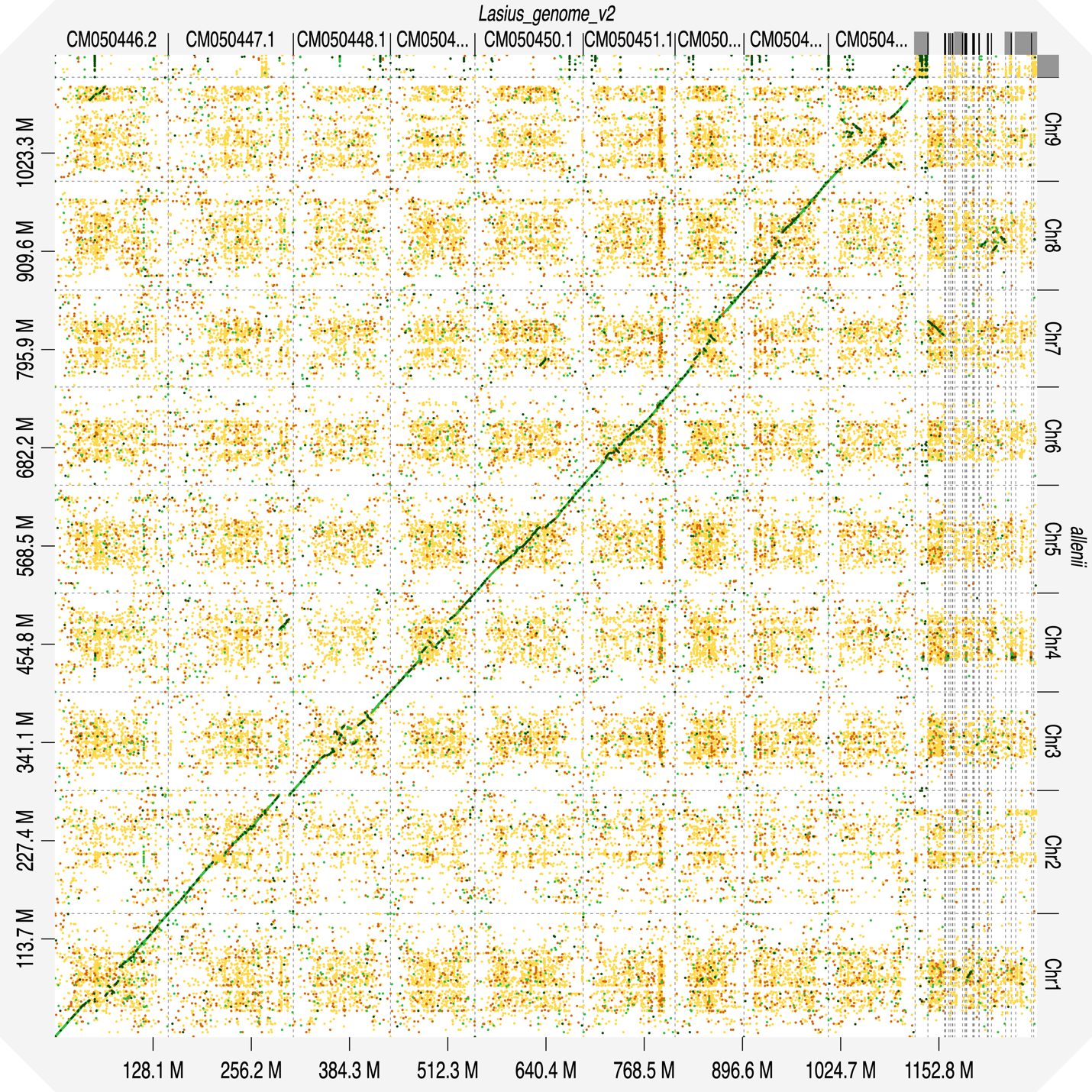

*Costus allenii* (Bee-pollinated)

*Costus lasius* (v2) (Hummingbird-pollinated)

Figure S3. Dotplot showing alignment between the *Costus lasius* genome assembly (x axis) and the *Costus allenii* genome assembly (y axis). Each dot represents a matching sequence with dark green representing the best matches. Dot plot generated with D-GENIES using Minimap2 alignment (<https://dgenies.toulouse.inra.fr/result/Costus_lasiusv2_vs_Costus_allenii>).

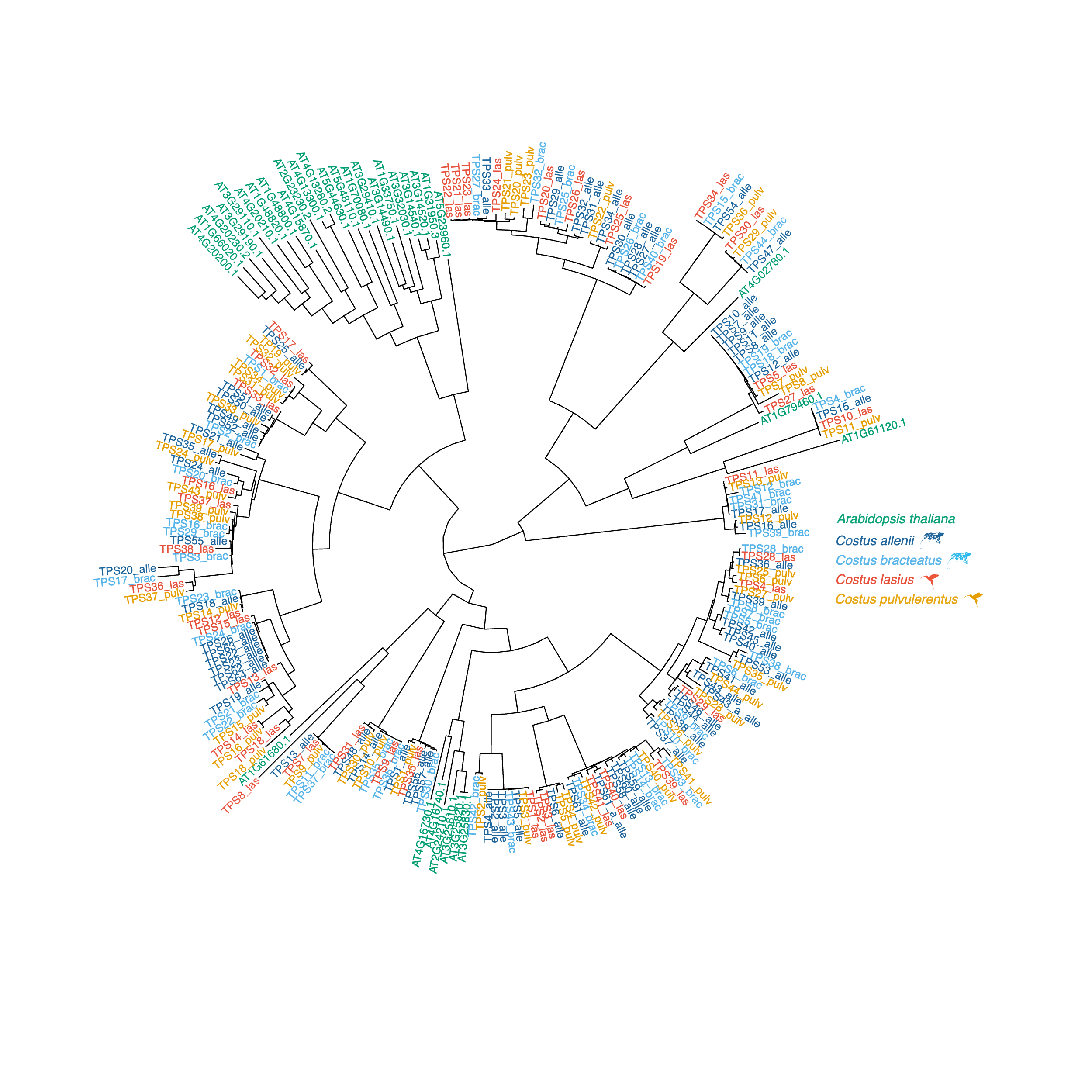

Figure S4. Phylogenetic analysis of terpene synthase genes. The phylogeny was constructed in IQ-TREE with 1000 bootstraps and model LG.

Table S1 Number of samples collected for each species. Tip labels are the same as those used in (Vargas et al. 2020).

| Species | Pollination group | Sample location | No. of samples |
| --- | --- | --- | --- |
| *Costus_allenii_98050* | Bee | UCSC greenhouse | 8 |
| *Costus_amazonicus_GUAL_18009* | Bee | UCSC greenhouse | 1 |
| *Costus_arabicus_98080* | Bee | UCSC greenhouse | 1 |
| *Costus_barbatus_19242* | Hummingbird | UCSC greenhouse | 1 |
| *Costus_bracteatus_18030* | Bee | UCSC greenhouse | 7 |
| *Costus_chartaceus_19269* | Hummingbird | UCSC greenhouse | 5 |
| *Costus claviger^+^* | Bee | UCSC greenhouse | 1 |
| *Costus_dubius_98074* | Bee | UCSC greenhouse | 3 |
| *Costus_erythrophyllus_98168* | Bee | UCSC greenhouse | 4 |
| *Costus_glaucus_98077* | Bee | La Gamba (Costa Rica) | 2 |
| *Costus_guanaiensis_var_macrostrobilus_18077* | Bee | UCSC greenhouse | 2 |
| *Costus_guanaiensis_var_tarmicus_19243* | Bee | UCSC greenhouse | 1 |
| *Costus_laevis_18032* | Bee | La Gamba, Las Cruces (Costa Rica) | 9 |
| *Costus_lasius_18080* | Hummingbird | UCSC greenhouse | 7 |
| *Costus_malortieanus_18036* | Bee | UCSC greenhouse | 7 |
| *Costus_montanus_19217* | Hummingbird | Monteverde (Costa Rica) | 2 |
| *Costus_osae_19005* | Hummingbird | UCSC greenhouse | 1 |
| *Costus phaeotrichus^+^* | Bee | UCSC greenhouse | 2 |
| *Costus_productus_98170* | Hummingbird | UCSC greenhouse | 5 |
| *Costus_pulverulentus_18042* | Hummingbird | UCSC greenhouse | 3 |
| *Costus pulverulentus x Costus scaber^+^* | Hummingbird | UCSC greenhouse | 1 |
| *Costus_scaber_18044** | Hummingbird | UCSC greenhouse | 1 |
| *Costus_scaber_18085** | Hummingbird | UCSC greenhouse | 2 |
| *Costus_spiralis_var_spiralis_18088* | Hummingbird | UCSC greenhouse | 1 |
| *Costus_stenophyllus_19227* | Hummingbird | La Gamba (Costa Rica) | 2 |
| *Costus_varzearum_19252* | Bee | UCSC greenhouse | 1 |
| *Costus_villosissimus_18090* | Bee | UCSC greenhouse | 7 |
| *Costus_vinosus_19273* | Bee | UCSC greenhouse | 4 |
| *Costus_wilsonii_18050* | Hummingbird | Las Cruces, Las Alturas (Costa Rica) | 8 |
| *Costus_woodsonii_18052* | Hummingbird | UCSC greenhouse | 1 |

**Costus scaber* is polyphyletic and so is represented by two tips on the phylogeny (Fig. 3).

^+^Not included in phylogenetic analyses

Table S2 Floral scent compounds identified in our dataset. Percentage of bee-pollinated and hummingbird-pollinated individuals in which each compound was detected. Adjusted (false discovery rate) chi-squared test p-values highlight the differences between bee- and hummingbird-pollinated samples. The identity of compounds in bold was confirmed with an authentic standard. For unidentified compounds, the most abundant m/z fragments are noted.

| Compound | Kovat’s RI | Type | Bee (%) | Hummingbird (%) | p-value |
| --- | --- | --- | --- | --- | --- |
| **alpha-pinene** | 931 | Monoterpene | 59 | 37 | NS |
| camphene | 944 | Monoterpene | 7 | 0 | NS |
| **sabinene** | 971 | Monoterpene | 14 | 0 | NS |
| **beta-pinene** | 973 | Monoterpene | 37 | 24 | NS |
| beta-myrcene | 991 | Monoterpene | 25 | 2 | NS |
| alpha-phellandrene | 1001 | Monoterpene | 8 | 0 | NS |
| Unknown (93, 121, 136) | 1014 | Monoterpene | 3 | 0 | NS |
| **cymene** | 1022 | Monoterpene | 34 | 10 | NS |
| **S-limonene** | 1026 | Monoterpene | 37 | 12 | NS |
| benzylalcohol | 1031 | Aromatic | 10 | 0 | NS |
| Unknown (79, 93, 105) | 1037 | Monoterpene | 12 | 0 | NS |
| **(*E*)-beta-ocimene** | 1048 | Monoterpene | 41 | 5 | <0.05 |
| gamma-terpinene | 1058 | Monoterpene | 15 | 0 | NS |
| **cineole** | 1078 | Monoterpene | 31 | 5 | NS |
| **terpinolene** | 1086 | Monoterpene | 3 | 0 | NS |
| methylbenzoate | 1095 | Aromatic | 17 | 0 | NS |
| **linalool** | 1099 | Monoterpene | 8 | 2 | NS |
| Unknown (53, 69, 79) | 1117 | Unknown | 42 | 20 | NS |
| Unknown (55, 69, 83) | 1176 | Unknown | 10 | 0 | NS |
| **methylsalicylate** | 1191 | Aromatic | 3 | 0 | NS |
| delta-elemene | 1338 | Sesquiterpene | 12 | 0 | NS |
| alpha-cubebene | 1350 | Sesquiterpene | 5 | 2 | NS |
| Unknown (91, 105, 161) | 1369 | Sesquiterpene | 10 | 2 | NS |
| alpha-copaene | 1376 | Sesquiterpene | 41 | 15 | NS |
| Unknown (91, 121, 161) | 1380 | Sesquiterpene | 12 | 0 | NS |
| **methylcinnamate** | 1382 | Aromatic | 7 | 0 | NS |
| Unknown (91, 108, 161) | 1390 | Sesquiterpene | 27 | 7 | NS |
| zingiberene | 1391 | Sesquiterpene | 3 | 0 | NS |
| Unknown (93, 105, 119) | 1401 | Sesquiterpene | 3 | 0 | NS |
| alpha-cedrene | 1412 | Sesquiterpene | 3 | 0 | NS |
| Unknown (93, 105, 119) | 1416 | Sesquiterpene | 15 | 2 | NS |
| (*E*)-beta-caryophyllene | 1419 | Sesquiterpene | 41 | 7 | <0.05 |
| gamma-elemene | 1434 | Sesquiterpene | 12 | 0 | NS |
| trans-bergamotene | 1437 | Sesquiterpene | 7 | 5 | NS |
| Unknown (69, 77, 93) | 1444 | Sesquiterpene | 15 | 5 | NS |
| beta-copaene | 1446 | Sesquiterpene | 5 | 0 | NS |
| alpha-humulene | 1454 | Sesquiterpene | 27 | 5 | NS |
| (*E*)-beta-farnesene | 1457 | Sesquiterpene | 8 | 0 | NS |
| amorphene | 1462 | Sesquiterpene | 12 | 0 | NS |
| (*Z*)-beta-Caryophyllene | 1468 | Sesquiterpene | 10 | 2 | NS |
| Unknown (105, 161, 204) | 1474 | Sesquiterpene | 5 | 0 | NS |
| Unknown (91, 133, 189) | 1476 | Sesquiterpene | 5 | 0 | NS |
| Unknown (91, 105, 161) | 1477 | Sesquiterpene | 3 | 5 | NS |
| germacrene D | 1481 | Sesquiterpene | 29 | 10 | NS |
| (*Z*)-beta-farnesene | 1485 | Sesquiterpene | 7 | 2 | NS |
| aciphyllene | 1495 | Sesquiterpene | 17 | 0 | NS |
| alpha-elemene | 1499 | Sesquiterpene | 19 | 5 | NS |
| Germacrene B | 1505 | Sesquiterpene | 17 | 2 | NS |
| alpha-farnesene | 1509 | Sesquiterpene | 20 | 5 | NS |
| gamma-cadinene | 1515 | Sesquiterpene | 24 | 5 | NS |
| delta-cadinene | 1524 | Sesquiterpene | 27 | 10 | NS |
| Unknown (91, 105, 161) | 1538 | Sesquiterpene | 5 | 0 | NS |
| elemol | 1558 | Sesquiterpene | 12 | 0 | NS |
| Unknown (69, 81, 95) | 1579 | Aromatic monoterpenoid | 10 | 0 | NS |
| caryophyllene oxide | 1583 | Sesquiterpene | 3 | 2 | NS |
| germacrene-d-4-ol | 1594 | Sesquiterpene | 7 | 2 | NS |
| Unknown (67, 93, 109) | 1610 | Unknown | 3 | 2 | NS |
| Unknown (105, 119, 161) | 1616 | Sesquiterpene | 15 | 2 | NS |
| Unknown (105, 161, 204) | 1641 | Sesquiterpene | 5 | 0 | NS |
| Unknown (93, 108, 126) | 1706 | Unknown | 3 | 0 | NS |

Table S3 BUSCO scores for transcriptomes searched against the Embryophyta database.

|  | *C. allenii* | *C. lasius* | *C. villosissimus* |
| --- | --- | --- | --- |
| Complete | 1401 (86.8%) | 1350 (83.7%) | 1362 (84.3%) |
| Complete and single copy | 430 (26.6%) | 590 (36.6%) | 372 (23.0%) |
| Complete and duplicated | 971 (60.2%) | 760 (47.1%) | 990 (61.3%) |
| Fragmented | 95 (5.9%) | 109 (6.8%) | 105 (6.5%) |
| Missing | 118 (7.3%) | 155 (9.5%) | 147 (9.2%) |
| Total searched | 1614 | 1614 | 1614 |

Table S4. *Costus allenii* genome assembly statistics

| N50 (Mb) | 120 |
| --- | --- |
| L50 | 5 |
| Total length (Gbp) | 1.137 |
| GC% | 40.07 |
| Number of Ns per 100 kbp | 0 |
| Longest contig/scaffold (Mbp) | 143 |
| Number of contigs/scaffolds | 423 |
| BUSCO complete | 98.8% (1594/1614) |
| BUSCO single copy | 90.8% (1465/1614) |

Table S5. BUSCO scores for high quality annotations of each *Costus* genome searched against the Embryophyta database.

|  | *C. allenii* | *C. bracteatus* | *C. lasius* | *C. pulverulentus* |
| --- | --- | --- | --- | --- |
| Complete | 1493 (92.5%) | 1457 (90.2%) | 1480 (91.7%) | 1466 (90.9%) |
| Complete and single copy | 1139 (70.6%) | 1116 (69.1%) | 1121 (69.5%) | 1152 (71.4%%) |
| Complete and duplicated | 354 (21.9%) | 341 (21.1%) | 359 (22.2%) | 314 (19.5%) |
| Fragmented | 12 (0.7%) | 25 (1.5%) | 19 (1.2%) | 21 (1.3%) |
| Missing | 109 (6.8%) | 132 (8.3%) | 115 (7.1%) | 127 (7.8%) |
| Total searched | 1614 | 1614 | 1614 | 1614 |
| Consistent lineage placement (OMArk) | 86% | 87% | 86% | 87% |
| Total gene number | 28,442 | 27,794 | 28,409 | 26,696 |

Table S6 Repetitive sequences identified in the *Costus allenii* genome

|  | Number of elements* | Length occupied (bp) | % of sequence |
| --- | --- | --- | --- |
| **Retroelements (Total)** | 288948 | 486993535 | **42.83** |
| SINEs | 2152 | 826805 | 0.07 |
| Total LINEs | 6762 | 4393918 | 0.39 |
| LINEs (L2/CR1/Rex) | 70 | 30823 | 0.00 |
| LINEs (L1/CIN4) | 6692 | 4363095 | 0.38 |
| Total LTRs | 280034 | 481772812 | 42.37 |
| LTR (BEL/Pao) | 73 | 27829 | 0.00 |
| LTR (Ty1/Copia) | 117108 | 306450247 | 26.95 |
| LTR (Gypsy/DIRS1) | 154657 | 169973381 | 14.95 |
| LTR (Retroviral) | 2402 | 1441219 | 0.13 |
| **DNA transposons** | 30178 | 15673277 | **1.38** |
| Hobo-Activator | 11839 | 6820577 | 0.60 |
| Tc1-IS630-Pogo | 979 | 173086 | 0.02 |
| MULE-MuDR | 5794 | 2944112 | 0.26 |
| Tourist/Harbinger | 8421 | 2765025 | 0.24 |
| **Rolling-circles** | 2113 | 1271374 | **0.11** |
| **Unclassified** | 735355 | 289141681 | **25.43** |
| **Total interspersed repeats** |  | 791808493 | **=69.64** |
| **Small RNA** | 7487 | 10666919 | 0.94 |
| **Simple repeats** | 121384 | 24304806 | 2.14 |
| **Low complexity** | 25736 | 1366253 | 0.12 |
| **Total repeats masked** |  | 829131231 | **72.92** |

Table S7 Repetitive sequences identified in the *Costus bracteatus* genome

|  | Number of elements* | Length occupied (bp) | % of sequence |
| --- | --- | --- | --- |
| **Retroelements (Total)** | 240419 | 343281137 | **35.81** |
| SINEs | 1006 | 549975 | 0.06 |
| Penelope | 135 | 38072 | 0.00 |
| Total LINEs | 5517 | 4802900 | 0.5 |
| LINEs (R1/LOA/Jockey) | 157 | 39875 | 0.00 |
| LINEs (L1/CIN4) | 5360 | 4763025 | 0.50 |
| Total LTRs | 233896 | 337928262 | 35.26 |
| LTR (BEL/Pao) | 309 | 204399 | 0.02 |
| LTR (Ty1/Copia) | 96965 | 178562625 | 18.63 |
| LTR (Gypsy/DIRS1) | 131382 | 154451766 | 16.11 |
| LTR (Retroviral) | 478 | 84941 | 0.01 |
| **DNA transposons** | 27071 | 15186655 | **1.58** |
| Hobo-Activator | 13283 | 7230193 | 0.75 |
| Tc1-IS630-Pogo | 1801 | 531912 | 0.06 |
| MULE-MuDR | 2533 | 2026276 | 0.21 |
| Tourist/Harbinger | 6077 | 2445486 | 0.26 |
| **Rolling-circles** | 3439 | 1822326 | **0.19** |
| **Unclassified** | 723175 | 294673790 | **30.74** |
| **Total interspersed repeats** |  | 653179654 | **=68.15** |
| **Small RNA** | 3049 | 2492596 | 0.26 |
| **Satellites** | 418 | 220408 | 0.02 |
| **Simple repeats** | 107332 | 8005154 | 0.84 |
| **Low complexity** | 21124 | 1103772 | 0.12 |
| **Total repeats masked** |  | 666577247 | **69.54** |

Table S8 Repetitive sequences identified in the *Costus lasius* genome

|  | Number of elements* | Length occupied (bp) | % of sequence |
| --- | --- | --- | --- |
| **Retroelements (Total)** | 353232 | 627944238 | **49.02** |
| SINEs | 246 | 231012 | 0.02 |
| LINEs (L1/CIN4) | 6311 | 4274994 | 0.33 |
| Total LTRs | 346675 | 623438232 | 48.67 |
| LTR (Ty1/Copia) | 156050 | 405840150 | 31.68 |
| LTR (Gypsy/DIRS1) | 183068 | 211184692 | 16.49 |
| LTR (Retroviral) | 1659 | 2779723 | 0.22 |
| **DNA transposons (Total)** | 26930 | 14928356 | **1.17** |
| hobo-Activator | 14915 | 8003790 | 0.62 |
| Tc1-IS630-Pogo | 943 | 459465 | 0.04 |
| MULE-MuDR | 1857 | 1892112 | 0.15 |
| Tourist-Harbinger | 5719 | 2103871 | 0.16 |
| **Rolling-circles** | 4743 | 1573269 | **0.12** |
| **Unclassified** | 726185 | 291748205 | **22.78** |
| **Total interspersed repeats** |  | 934620799 | **=72.97** |
| **Small RNA** | 5707 | 16921341 | 1.32 |
| **Satellites** | 1073 | 100316 | 0.01 |
| **Simple repeats** | 121862 | 17247730 | 1.35 |
| **Low complexity** | 23049 | 1239249 | 0.10 |
| **Total repeats masked** |  | 971471692 | **75.84** |

Table S9 Repetitive sequences identified in the *Costus pulverulentus* genome

|  | Number of elements* | Length occupied (bp) | % of sequence |
| --- | --- | --- | --- |
| **Retroelements (Total)** | 215251 | 199407364 | **27.10** |
| SINEs | 1595 | 550790 | 0.07 |
| Total LINEs | 7594 | 4350090 | 0.59 |
| LINEs (RTE/Bov-B) | 171 | 55791 | 0.01 |
| LINEs (L1/CIN4) | 7303 | 4268153 | 0.58 |
| Total LTRs | 206062 | 194506484 | 26.43 |
| LTR (BEL/Pao) | 2693 | 786971 | 0.11 |
| LTR (Ty1/Copia) | 77676 | 75627603 | 10.28 |
| LTR (Gypsy/DIRS1) | 120490 | 115423253 | 15.69 |
| LTR (Retroviral) | 1508 | 441970 | 0.06 |
| **DNA transposons (Total)** | 22872 | 11781701 | **1.60** |
| hobo-Activator | 10694 | 5108156 | 0.69 |
| Tc1-IS630-Pogo | 1113 | 254037 | 0.03 |
| MULE-MuDR | 2803 | 2184543 | 0.30 |
| Tourist-Harbinger | 5475 | 2028061 | 0.28 |
| **Rolling-circles** | 982 | 476438 | **0.06** |
| **Unclassified** | 734178 | 245218168 | **33.32** |
| **Total interspersed repeats** |  | 456407233 | **=62.02** |
| **Small RNA** | 3364 | 793375 | 0.11 |
| **Satellites** | 81 | 35422 | 0.00 |
| **Simple repeats** | 98952 | 5596530 | 0.76 |
| **Low complexity** | 18654 | 990059 | 0.13 |
| **Total repeats masked** |  | 463924390 | **63.05** |

Table S10 All gene ontology (GO) terms that are significantly enriched in differentially expressed genes. Gene sets were analyzed separately for each pairwise comparison and for genes upregulated in each species. weightFisher=Fisher exact p-value, BP=Biological Process, MF=Molecular Function, CC=Cellular Component.

| Comparison | Upregulated | GO ID | Term | weightFisher | Type |
| --- | --- | --- | --- | --- | --- |
| *C. allenii* vs *C. lasius* | *C. allenii* | GO:0043167 | Ion binding | 0.034 | MF |
| *C. allenii* vs *C. lasius* | *C. allenii* | GO:0008168 | Methyltransferase activity | 0.034 | MF |
| *C. allenii* vs *C. lasius* | *C. lasius* | GO:0006950 | Response to stress | 0.022 | BP |
| *C. allenii* vs *C. lasius* | *C. lasius* | GO:0046872 | Metal ion binding | 0.028 | MF |
| *C. allenii* vs *C. lasius* | *C. lasius* | GO:0016791 | Phosphatase activity | 0.034 | MF |
| *C. allenii* vs *C. lasius* | *C. lasius* | GO:0035251 | UDP-glucosyltrasnferase activity | 0.034 | MF |
| *C. allenii* vs *C. lasius* | *C. lasius* | GO:0016020 | Membrane | 0.038 | CC |
| *C. allenii* vs *C. villosissimus* | *C. allenii* | GO:1900150 | Regulation of defense response to fungus | 0.0053 | BP |
| *C. allenii* vs *C. villosissimus* | *C. allenii* | GO:0042221 | Response to chemical | 0.0305 | BP |
| *C. allenii* vs *C. villosissimus* | *C. allenii* | GO:0016747 | Acyltransferase activity | 0.015 | MF |
| *C. allenii* vs *C. villosissimus* | *C. allenii* | GO:0043169 | Cation binding | 0.017 | MF |
| *C. allenii* vs *C. villosissimus* | *C. allenii* | GO:0000166 | Nucleotide binding | 0.035 | MF |
| *C. allenii* vs *C. villosissimus* | *C. allenii* | GO:0016787 | Hydrolase activity | 0.048 | MF |
| *C. allenii* vs *C. villosissimus* | *C. villosissimus* | GO:0010264 | Myo-inositol hexakisphosphate biosynthetic process | 0.0087 | BP |
| *C. allenii* vs *C. villosissimus* | *C. villosissimus* | GO:0035251 | UDP-glucosyltrasnferase activity | 0.0049 | MF |
| *C. allenii* vs *C. villosissimus* | *C. villosissimus* | GO:0050285 | Sinapine esterase activity | 0.0014 | MF |
| *C. allenii* vs *C. villosissimus* | *C. villosissimus* | GO:0003824 | Catalytic activity | 0.0467 | MF |
| *C. lasius* vs *C. villosissimus* | *C. lasius* | GO:0048731 | System development | 0.024 | BP |
| *C. lasius* vs *C. villosissimus* | *C. lasius* | GO:0042221 | Response to chemical | 0.038 | BP |
| *C. lasius* vs *C. villosissimus* | *C. lasius* | GO:0046872 | Metal ion binding | 0.0089 | MF |
| *C. lasius* vs *C. villosissimus* | *C. lasius* | GO:0043169 | Cation binding | 0.0183 | MF |
| *C. lasius* vs *C. villosissimus* | *C. lasius* | GO:0004857 | Enzyme inhibitor activity | 0.0239 | MF |
| *C. lasius* vs *C. villosissimus* | *C. lasius* | GO:0000166 | Nucleotide binding | 0.0428 | MF |
| *C. lasius* vs *C. villosissimus* | *C. lasius* | GO:0005576 | Extracellular region | 0.005 | CC |
| *C. lasius* vs *C. villosissimus* | *C. villosissimus* | GO:0035251 | UDP-glucosyltrasnferase activity | 0.027 | MF |
